# Intratumor Heterogeneity and Circulating Tumor Cell Clusters

**DOI:** 10.1101/113480

**Authors:** Zafarali Ahmed, Simon Gravel

**Affiliations:** Department of Biology, McGill University, Montreal, Quebec, Canada; Department of Human Genetics, McGill University, Montreal, Quebec, Canada

## Abstract

Genetic diversity plays a central role in tumor progression, metastasis, and resistance to treatment. Experiments are shedding light on this diversity at ever finer scales, but interpretation is challenging. Using recent progress in numerical models, we simulate macroscopic tumors to investigate the interplay between growth dynamics, microscopic composition, and circulating tumor cell cluster diversity. We find that modest differences in growth parameters can profoundly change microscopic diversity. Simple outwards expansion leads to spatially segregated clones and low diversity, as expected. However, a modest cell turnover can result in an increased number of divisions and mixing among clones resulting in increased microscopic diversity in the tumor core. Using simulations to estimate power to detect such spatial trends, we find that multiregion sequencing data from contemporary studies is marginally powered to detect the predicted effects. Slightly larger samples, improved detection of rare variants, or sequencing of smaller biopsies or circulating tumor cell clusters would allow one to distinguish between leading models of tumor evolution. The genetic composition of circulating tumor cell clusters, which can be obtained from non-invasive blood draws, is therefore informative about tumor evolution and its metastatic potential.

**Highlights:**

1. Numerical and theoretical models show interaction of front expansion, mutation, and clonal mixing in shaping tumor heterogeneity.
2. Cell turnover increases intratumor heterogeneity.
3. Simulated circulating tumor cell clusters and microbiopsies exhibit substantial diversity with strong spatial trends.
4. Simulations suggest attainable sampling schemes able to distinguish between prevalent tumor growth models.

## Introduction

Most cancer deaths are due to metastasis of the primary tumor, which complicates treatment and promotes relapse (Holohan et al. 2013; Vanharanta and Massagu´e 2013; Quail and Joyce 2013; Steeg 2016). Circulating tumor cells (CTC) are bloodborne enablers of metastasis that were first detected in the blood of patients after death (Ashworth 1869) and can now be captured using a variety of devices (Joosse, Gorges, and Pantel 2014; Sarioglu et al. 2015; Glynn et al. 2015; Siravegna et al. 2017) allowing us to study their origins and implications for metastasis (Massagu´e and Obenauf 2016; Lambert, Pattabiraman, and Weinberg 2017). Counts of single CTCs have been used to predict tumor progression (Cristofanilli et al. 2005; Krebs, Sloane, et al. 2011; Siravegna et al. 2017) and monitor curative and palliative therapies in a vast array of cancer types (D. Hayes et al. 2002; Wülfing et al. 2006; Aceto, Toner, et al. 2015; Siravegna et al. 2017). CTCs have also been isolated in clusters of up to 100 cells (Marrinucci et al. 2012; Aceto, Bardia, et al. 2014; Glynn et al. 2015; Au et al. 2017). These CTC clusters, though rare, are associated with more aggressive metastatic cancer and poorer survival rates in mice and breast and prostate cancer patients (Liotta, Kleinerman, and Saldel 1976; Glaves 1983; Aceto, Bardia, et al. 2014; Cheung et al. 2016). Cellular growth within tumors follows Darwinian evolution with sequential accumulation of mutations and selection resulting in subclones of different fitness (Nowell 1976; Burrell et al. 2013; Williams et al. 2016). Certain classes of mutations are known to give cancer cells advantages beyond local growth rates. For example, acquiring mutations in *ANGPTL4* in breast tumors does not appear to provide a growth advantage to cells in the primary, however it enhances metastatic potential to the lungs (Padua et al. 2008). Similarly, breast tumors are more likely to metastasize into the lung or brain if they acquire mutations in *TGFβ* or *ST6GALNAC5*, respectively (Padua et al. 2008; Bos et al. 2009; Peinado et al. 2017). These genes are referred to as metastasis progression genes or metastasis virulence genes (Lorusso and Rüegg 2012).

Mutations, including those in metastasis progression and virulence genes, are not uniformly distributed in the tumor. Tumors show substantial intratumoral heterogeneity (ITH) (Navin et al. 2010; Gerlinger, Rowan, et al. 2012; Sottoriva et al. 2015; McGranahan and Swanton 2015; McGranahan and Swanton 2017) where subclones have private mutations that can lead to subclonal phenotypes (J. Zhang et al. 2014; Gerlinger, Horswell, et al. 2014; Yates et al. 2015; Morrissy et al. 2017; Peinado et al. 2017). A high degree of ITH can allow tumors to explore a wide range of phenotypes relevant to metastatic outlook. Additionally, ITH can contribute to therapy resistance and relapse (Holohan et al. 2013; Hiley et al. 2014). Studying ITH is therefore important for understanding cancer progression and improving therapeutic and prognostic decisions (Hiley et al. 2014; Jamal-Hanjani, Hackshaw, et al. 2014; Alizadeh et al. 2015; Andor et al. 2016). To capture the complete mutational spectrum of a primary tumor, multiple study designs have been proposed that divide the tumor into regionally representative samples, known as multiregion sequencing (Gerlinger, Rowan, et al. 2012; Gerlinger, Horswell, et al. 2014; J. Zhang et al. 2014; Yates et al. 2015; Ling et al. 2015).

Next-generation sequencing (NGS) and molecular profiling has shown that CTCs have similar genetic composition to both the primary and metastatic lesions (Heitzer et al. 2013; Brouwer et al. 2016; Siravegna et al. 2017). Sequencing of CTCs can therefore be used as a non-invasive liquid biopsy to study tumors and tumor heterogeneity, monitor response to therapy, and determine patient-specific course of treatment (Powell et al. 2012; Heitzer et al. 2013; Krebs, Metcalf, et al. 2014; Hodgkinson et al. 2014).

Here we use simulations to assess whether genetic heterogeneity within individual circulating tumor cell clusters can be informative about solid tumor progression. Because CTC clusters are thought to originate from neighboring cells in the tumor (Aceto, Bardia, et al. 2014), heterogeneity within CTC clusters is closely related to cellular-scale genetic heterogeneity within tumors. We therefore interpret our simulation results as informative about both micro-biopsies and circulating tumor cell clusters.

We used an extension^1^ of the simulator described in Waclaw *et al.* (2015) to study the interplay of tumor dynamics, CTC cluster diversity, and metastatic outlook. We first consider tumor-wide heterogeneity patterns, and find that the overall distribution of common allele frequencies is well described by a recent analytical model of tumor growth (Fusco et al. 2016) that assumes neutrality and no turnover: the global patterns of diversity are relatively robust to modest departures from these assumptions. We then show that fine-scale tumor heterogeneity, and therefore CTC cluster composition, depend more sensitively on the turnover dynamics of the tumor. We discuss consequences for metastatic outlook and, by simulating multiregion sequencing studies of micro-biopsies, show that currently achievable sample sizes would be well powered to identify spatial trends and distinguish between leading models of tumor evolution.

## Simulation Model

To simulate the growth of solid tumors, we use TumorSimulator^2^ (Waclaw et al. 2015). The software is able to simulate a tumor containing 10^8^ cells, or roughly 1 cubic centimeter (Del Monte 2009), in less than 24 core-hours. The tumor consists of cells that occupy points on a 3D lattice. Cells do not move in this model: The tumor evolves through cell division and death.

Empty lattice sites are assumed to contain normal cells which are not modelled in TumorSimulator.

Each cell has an associated list of genetic alterations which represent single nucleotide polymorphisms (SNPs) that can be either passenger or driver. Driver mutations increase the growth rate by a factor 1 + *s*, where *s ≥*0 is the average selective advantage of a driver mutation.

The simulation begins with a single cell that already has an unlimited growth potential. Tumor growth then proceeds by selecting a mother cell randomly. It then divides with a probability proportional to *b*_0_(1 + *s*)^*k*^ (rescaled by the maximal birth rate of all cells in the tumor, such that this probability is ≤ 1) where *b*_0_ is the inital birth rate and *k* is the number of driver mutations in that cell. New cells are given new passenger and driver mutations according to two independent Poisson distributions parameterized by haploid mutation rates *µ_p_* and *µ_d_* so that the maximal frequency in a tumor is one. The mother cell dies with a probability proportional to the death rate *d* (rescaled in a similar manner as the birth rate), independent of whether it succesfully reproduced. The simulation ends when there are 10^8^ cells in the tumor, unless otherwise specified. To facilitate comparison, we first set parameters *b*_0_*, s, µ_p_*, and *µ_d_* to match those used in Waclaw *et al.* (2015). When comparing to experimental data in Ling *et al.* (2015), we adjust the passenger mutation rate to match empirical observations (See further details of the algorithm and complete description of parameters in Supplemental Information and Table S2 respectively).

We consider three turnover scenarios corresponding to three models for the death rate *d*: (i) No turnover (*d* = 0), corresponding to simple clonal growth (Hallatschek et al. 2007; Fusco et al. 2016); (ii) Surface Turnover (*d*(*x, y, z*) *>* 0 only if *x, y, z* is on the surface), corresponding to a quiescent core model (Shweiki et al. 1995) (iii) Turnover (*d >* 0 everywhere), a model favored in Waclaw et al. 2015 to explore ITH.

## Results

### Global composition

To determine the effect of the growth dynamics on global intratumor heterogeneity, we first consider the distribution of allele frequencies (or allele frequency spectra, AFS) for different turnover models (Fig 1, S1). In all cases, a majority of driver and passenger genetic variants are at frequency less than 1%, as expected from theoretical and empirical observations (e.g., Wang et al. 2014; Fusco et al. 2016). Passenger mutations represent the bulk of ITH independently of the selection coefficient (Fig S2), consistent with the theoretical and experimental evidence that neutral evolution drives most ITH (Williams et al. 2016). For simulations with low to moderate death rate, *d ∈* {0.05, 0.1, 0.2} and *s* = 1%, we find that the frequency spectra are very similar across the three turnover models (Fig 1, S1, S2): A low death rate has little impact on the global composition of a tumor.

**Figure 1:**
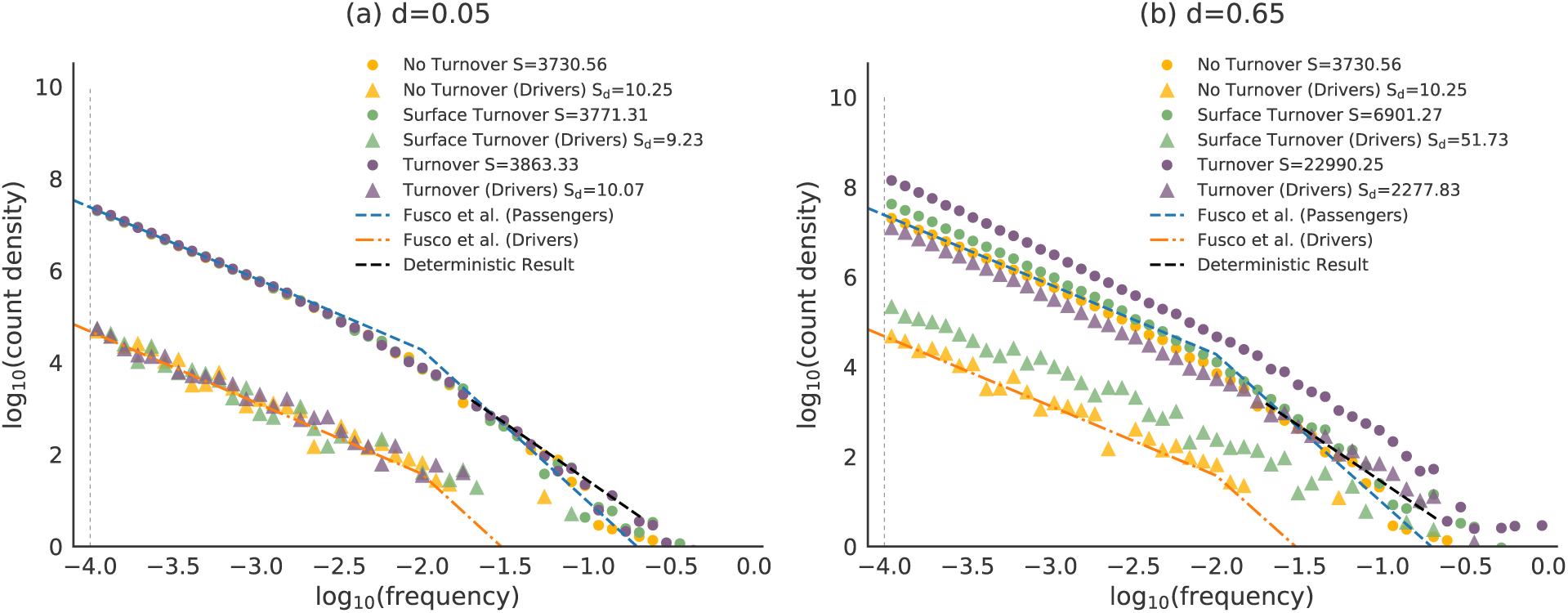
Frequency Spectra for the Primary Tumor at (a) low death rate and (b) high death rate for all mutations (circles) and driver mutations (triangles). At low death rate, the frequency spectra are nearly indistinguishable, whereas for higher death rate, the turnover model produces elevated diversity across the frequency spectrum for both driver and neutral mutations. The total number of somatic mutations, *S*, and the total number of driver mutations, *S_d_*, in the tumor is shown in the legend (average of 15 simulations). The vertical gray dotted line shows the minimum frequency of mutations returned by TumorSimulator. The black dotted line shows the asymptotic result of a geometric model with a scaling of *ζ* = 30 and is described in Supplementary Section S.5. The blue and oranged dashed lines shows the result from Fusco *et al.*. Fig S1 and S2 show simulations with intermediate values of *d* and different values of *s*.

When the death rate is increased to *d* = 0.65, as in Waclaw *et al.* (2015), the different models produce distinct frequency spectra (Fig 1b). Waclaw *et al* (2015) considered the number of high-frequency driver mutations as a measure of diversity, which is a simple summary statistic of the AFS. As in Waclaw *et al.*, we find that the number of high-frequency drivers is higher in the turnover model than in the no turnover model. Waclaw *et al.* interpreted this observation as an indication that turnover reduces diversity, because high frequencies suggest a larger number of dominant clones. However, we find that diversity, as measured by the number of polymorphic sites, is in fact increased for all types of variants and at all frequencies. The number of somatic mutations in the turnover model is 3.4 times higher than in the surface turnover model and 6.2 times higher than in the no turnover model. This is primarily due to a higher number of cell divisions required to reach a given tumor size when cell death occurs throughout the tumor (Table S1). The Waclaw *et al*. model uses a death rate of *d* = 0.65, which is a staggering 95% of the birth rate. The turnover model therefore has 8.3 times more cell divisions to reach a given size, and the surface turnover has 4 times more cell divisions than the no turnover model (Table S1).

Fig 1a exhibits two distinct power-law behaviors, a high-frequency power-law distribution *ϕ*(*f*) of mutations with frequency *f* scaling as *ϕ*(*f*) ~ f ^−2.5^, and a low-frequency scaling as *ϕ*(*f*) ~ f ^−1.61^. This scaling is present in the neutral case with no turnover (Fig S2a). Scaling laws in the distribution of allele frequencies have attracted considerable interest, harking back to the Wright-Fisher model for a constant-sized population (the “standard neutral model”) which predicts *ϕ*(*f*) ~ f ^−1^(Wright 1931; R. A. Fisher 1999). Population growth leads to an excess of rare variants: Tumor models that account for exponential population growth in a coalescent or branching process framework (Ohtsuki and Innan 2017) predict *ϕ*(*f*) *f ^−^*^1^ to *ϕ*(*f*) *f ^−^*^2^, depending on model parameters. A more directly applicable theoretical model was developed in Fusco *et al.* (2016) to model outwards growth of a bacterial colony or tumor, without turnover. Based on experimental and simulation data, also showing two scaling regimes, Fusco *et al.* considered a low-frequency regime containing “bubbles” (mutations that are cut off from the surface) and a high-frequency regime consisting of “sectors” (mutations that kept on with surface growth). They then used a KardarParisi-Zhang model (Kardar, Parisi, and Y.-C. Zhang 1986) of surface growth that predicts scaling laws of *ϕ*(*f*) ~ f ^−1.55^ at low frequencies, and of *ϕ*(*f*) ~ f ^−3.3^ at high frequencies (assuming a rough tumor surface). Supplementary Section S.5 also provides a simplified deterministic and neutral geometric model for sectors which predicts a decay for common variants *ϕ*(*f*) ~ f ^−2.5^ (Figs 1 and S2).

We adapted the continuity matching from Fusco *et al.* for distributions of allele frequencies (Fig S3), leading to predicted transition at frequency *f_c_* = 10^*−*1.7^. Both scaling laws and transition point are in excellent agreement with observations, with no fitting parameters (Fig 1). However some departures are visible at extremely low frequencies (Fig S4a).

Even though the Fusco et al. model assumes no turnover, it is relatively robust to modest turnover. For *d* = 0.2, there is a 20% increase of the overall number of segregating sites, but no difference in the overall scaling of common variants (Fig S2). Even in the large turnover regime (*d* = 0.65), the two distinct scaling laws are still clearly visible, suggesting that the distinction between bubbles and sectors is a useful construct despite the massive turnover. Similarly, selection has a weak effect on global patterns of passenger diversity except under the presence of extremely strong turnover (Fig S2). Turnover does increase the discrepancy between simulations and the Fusco et al. model for very rare variants (Fig S4). Supplementary section S.9 presents an extension to the Fusco et al model that accounts for the role of cell turnover in increasing the number of mutations in the tumor core (Fig. S4b).

### Cluster diversity depends on sampling position and turnover rate

To study the effect of cluster size, position of origin, and evolutionary model on CTC cluster composition, we sampled groups of cells across tumors (More details in Supplementary Section ‘CTC cluster synthesis’). To assess genetic heterogeneity within clusters, we consider the number of distinct somatic mutations, *S*(*n*), among cells in clusters of size *n*.

As expected, we find that larger CTC clusters have more somatic mutations (Fig 2, S5). Whereas moderate turnover had little impact on the tumor-wide number or frequency distribution of segregating sites, it can lead to a 5-fold increase in the number of segregating sites observed in small clusters: Clusters from models with low turnover have many more somatic mutations than in the no turnover model (Fig 2a,b). Surface turnover with low death rates *d ∈* {0, 0.05, 0.1, 0.2} has little effect on cluster diversity (Fig S5).

**Figure 2:**
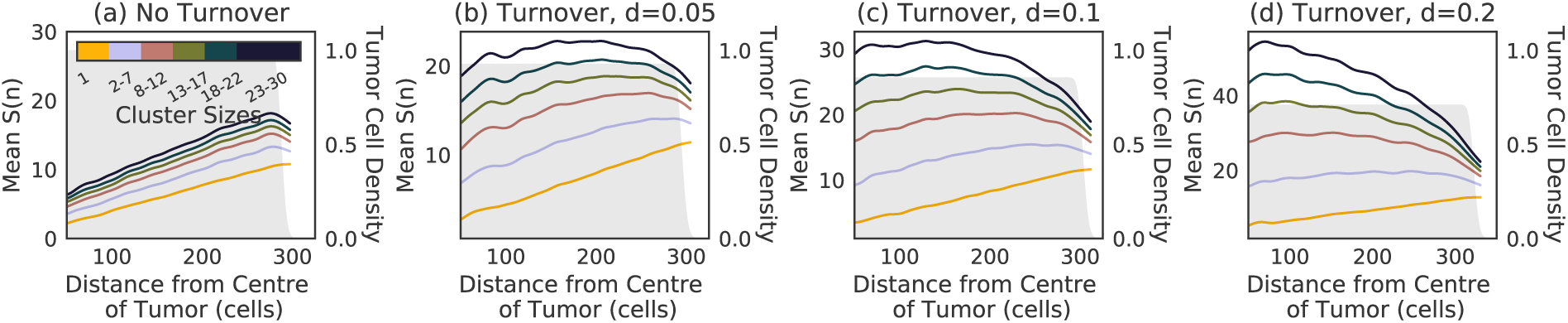
**Number of somatic mutations per cluster** as a function of cluster size and position for a model with death rate set to (a) *d* = 0 (no turnover) (b) *d* = 0.05, (c) *d* = 0.1 and (d) *d* = 0.2. The number of mutations in single CTCs increases at the edge, reflecting the larger number of cell divisions. The trend is reversed for larger clusters with at higher death rate. The shaded gray area represents the density of tumor cells at each position. The smoothed curves were obtained by a Gaussian weighted average using weight *w_i_*(*x*) = exp(*−*(*x − x_i_*)^2^), where *x_i_* is the distance from the centre of the tumor. See Fig S5 and S6 for the surface turnover model and turnover model with *d* = 0.65 respectively.

Fig 2 also shows the relationship between a CTC cluster’s shedding location (i.e. its distance to the tumor center-of-mass when it was sampled) and its genetic content. No turnover and surface turnover models show similar trends of increasing diversity with distance (Fig S5). Full turnover models show an opposite trend of decreasing diversity with distance in clusters of intermediate size (Fig 2b-d and S6 for *d* = 0.1, 0.2, and 0.65, respectively).

The number of distinct somatic mutations per cluster *S*(*n*) shows a dip near the tumor surface where the cell density has not yet reached equilibrium (Fig 2 and S6). This is the result of two transient effects. First, the earliest cells to populate the expansion front have experienced fewer divisions than the later cells, thus the average number of mutations in cells at a given distance from the tumor center increases as the front progresses. Second, the cells that first populate empty areas in the expansion front are more closely related to each other: If a cell has only one neighbor, it must descend directly from that neighbor; if a cell has 26 neighbors, it only has a 1/26 chance of descending directly from any given immediate neighbor — the time to the most recent common ancestor between neighbors increases as space fills up. Fig S8, which shows how *S*(*n*) changes as the tumor expands from size 10^6^ to 10^8^, also shows that this dip travels with the expansion front.

Fig S8 also shows how *S*(*n*) changes within the core of the tumor as it expands to eventually generate the patterns seen in Fig 2. Two processes increase cluster diversity within the core: new mutations and mixing among existing clones. To disentangle the effect of these two processes, we produce an equivalent time-course simulation where new mutations are turned off when the tumor reaches 10^6^ cells, leaving only clone mixing to increase genetic diversity. Fig S9 shows contrasting effects in the core and edge of the tumor: the diversity in edge clusters decreases over time because of serial founder effects. By contrast, the number of somatic mutations in clusters near the centre of the tumor increases: Mixing causes an increase in the number of distinct somatic mutations present in a cluster of a given size by bringing together cells from more distant backgrounds, increasing the effective population size. This leads to a roughly linear increase of cluster diversity with distance from the tumor edge. For *d* = 0.1 and clusters of 20 cells, the number of somatic mutations at the tumor centre increases from 5 to 8 as the tumor grows from 10^6^ to 10^8^ cells (Fig S9). The number of somatic mutations further increases to 13 if mutations are allowed in the core of the tumor (Fig S8): new mutations in this case contribute more to diversity in the core than clonal mixing.

Fig S10 show an alternate representation of this effect: we visualize the coalescence trees for neighbourhoods of 30 cells at the center and edges of the tumor. Neighbourhoods near the center of the tumor have longer terminal branches as there was more time for additional mutations to accumulate. This effect is particularly pronounced as the death rate increases. Neighbourhoods near the edge share a larger proportion of the trunk indicating that the cells have a recent common ancestor as a consequence of the serial founder effect: the height of the trees are higher at the edge, but the sum of branch lengths (i.e., *S*(*n*)) are higher in the center for the turnover model.

### Comparison with multi-region sequencing data

We did not have access to large-scale sequencing data for micro-biopsies. To illustrate predictions of our model, we therefore used multi-region sequencing data from a Hepatocellular Carcinoma (HCC) patient presented in Ling *et al.* (2015) (Fig 3a). The HCC data contained 23 sequenced samples from a single tumor each with ≈ 20, 000 cells. We therefore used our sampling scheme to simulate 23 biopsies of comparable sizes (20, 000 cells). The distance measurements were made using ImageJ (Schneider, Rasband, and Eliceiri 2012) and Fig S1 from Ling et al. 2015. Since Ling *et al.* (2015) could only reliably call variants at more than 10% frequency, we used a similar frequency cutoff in our simulations. The HCC data does not show a clear spatial trend (Fig 3a) whereas simulations with and without turnover had detectable trends at comparable sample size (Fig 3c,d).

**Figure 3:**
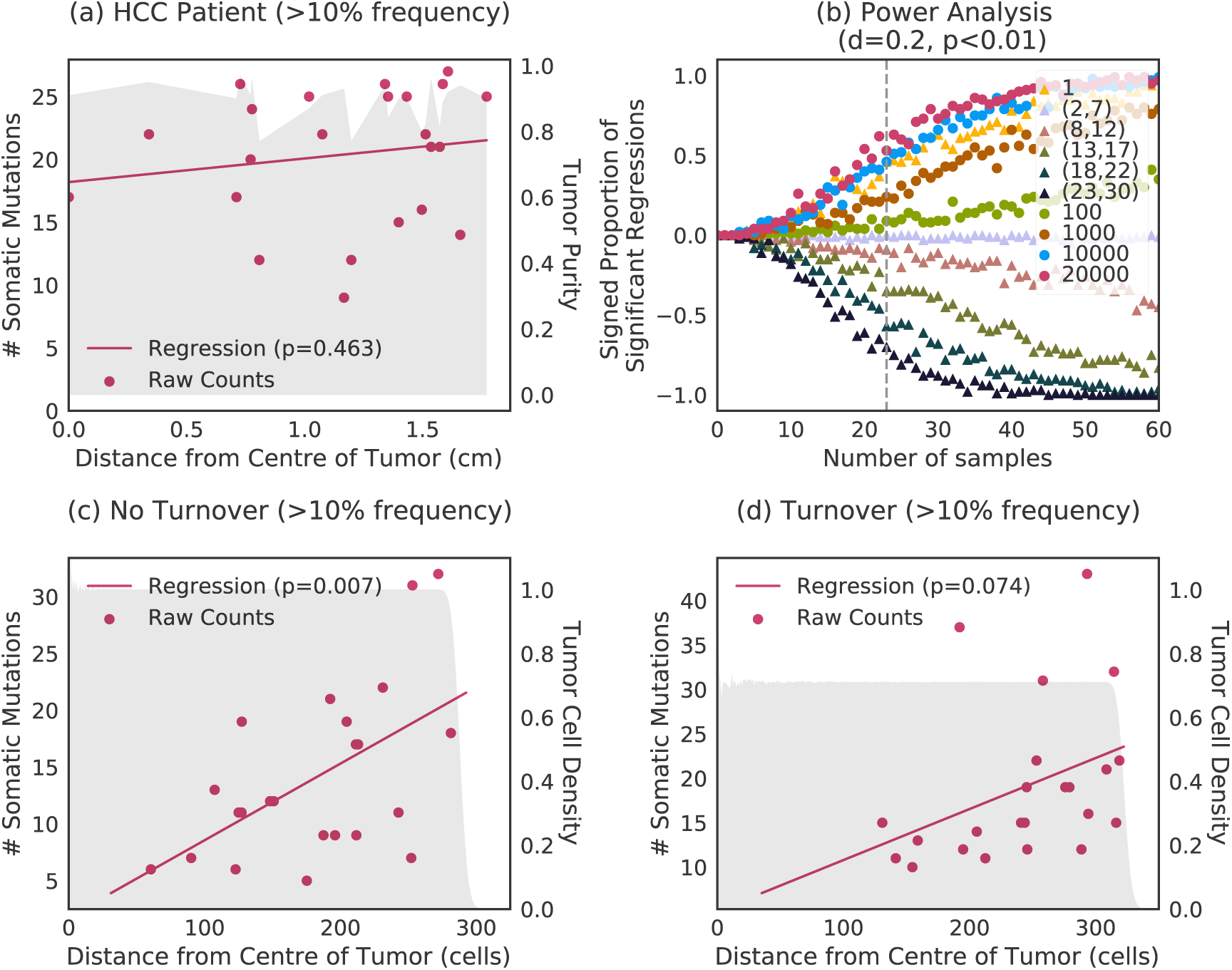
Comparison of simulated multi-region NGS with empirical hepatocellular carcinoma. (a) Spatial distribution and regression of the number of somatic mutations of 23 samples (20,000 cells each) in hepatocellular carcinoma patient. (b) Power to identify spatial trends in diversity as a function of cluster size and sample size (biopsies with over 100 cells have a frequency cutoff of *>* 10%, while smaller clusters have no frequency cutoff). The signed proportion of significant regressions counts the number of regressions that were significant (*p <* 0.01) for positive and negative slopes (See Supplementary Section S.3). Spatial trends in simulated tumors with sampling schemes as in (a), without turnover (c) and with turnover (d). The shaded gray area of (a) represents the tumor purity of the samples at each position. The shaded gray area of (c) and (d) represents the density of tumor cells at each position. See also Fig S11 for power analyses for the no turnover and different cell death rats *d*.

We therefore investigated the study design that would be needed to effectively distinguish between the different models proposed here. Based on simulations, power depends on cluster size, number of clusters sampled, and the choice of frequency cutoff (Fig 3b and S11). For a sample of 23 biopsies with ≈ 20, 000 cells each and a frequency cutoff of *>* 10%, we only have 50% power to detect a spatial trend in both turnover and no turnover models (Fig S11).

Spatial trends observed in Figs 2 and S5 are barely detectable with the current sample size but could be detected with modest increases in sample size or decreases in the frequency cutoff (Fig 3b). The choice of frequency cutoff can qualitatively affect spatial trends. Biopsies containing tens of thousands of cells with a 10% frequency cutoff show an increase in diversity at the edge of the tumor across all turnover models, with the number of spatially distributed samples needed to detect the trend reliably close to 40, roughly twice the size of the HCC dataset. If all mutations could be reliably detected, including at frequencies below 1%, spatial patterns should be apparent with only 10 biopsies, and these would highlight qualitative differences between the models, with increased diversity in the core for turnover models (Fig S11).

Small cluster sequencing, by focusing on globally rare but locally common variation, easily captures such differences in growth models. Approximately 30 deep sequenced small cluster (23-30 cells) samples are sufficient to reliably reveal qualitative difference between turnover models that neither single cells nor large biopsies capture, even at low (1%) frequency cutoffs (Fig S11).

### CTC clusters derived from turnover models are more likely to contain virulent mutations

Metastasis is an inefficient process (Massagu´e and Obenauf 2016) in that most CTCs are eliminated from the circulatory system or fail to survive in the new microenvironment. We hypothesize that the genetic composition of CTC clusters influences the likelihood of implantation into a new microenvironment. More specifically, genetic heterogeneity within a cluster may contribute to implantation by increasing the likelihood that a metastasis progression mutation is present. If a cluster has *S* somatic mutations, and each mutation has a small probability *p ≪* 1 of being a metastasis progression or virulence gene, the probability of having at least one such metastasis virulence gene is 1 − (1 − p)^*S*^ ≈Sp.

Diverse CTC clusters do not carry more virulent mutations, on average, than homogeneous ones, but they are more likely to carry *some* virulent mutations because of the increased diversity. Unless implantation probability is exactly proportional to the number of cells carrying virulent mutations in a cluster, which seems unlikely, diversity will impact implantation rate.

To compare the increased likelihood that CTC clusters possess metastatic progression genes compared to single CTCs, we determine the relative increase in the number of distinct somatic mutations in a CTC cluster versus a single CTC and refer to it as the *cluster advantage*, *A*(*n*). To disentangle the contributions from the microscopic and macroscopic diversity, as well as cluster size effects, we compute the cluster advantage for clusters composed of neighboring cells, as well as for random sets of cells sampled across the tumor (Fig 4).

**Figure 4:**
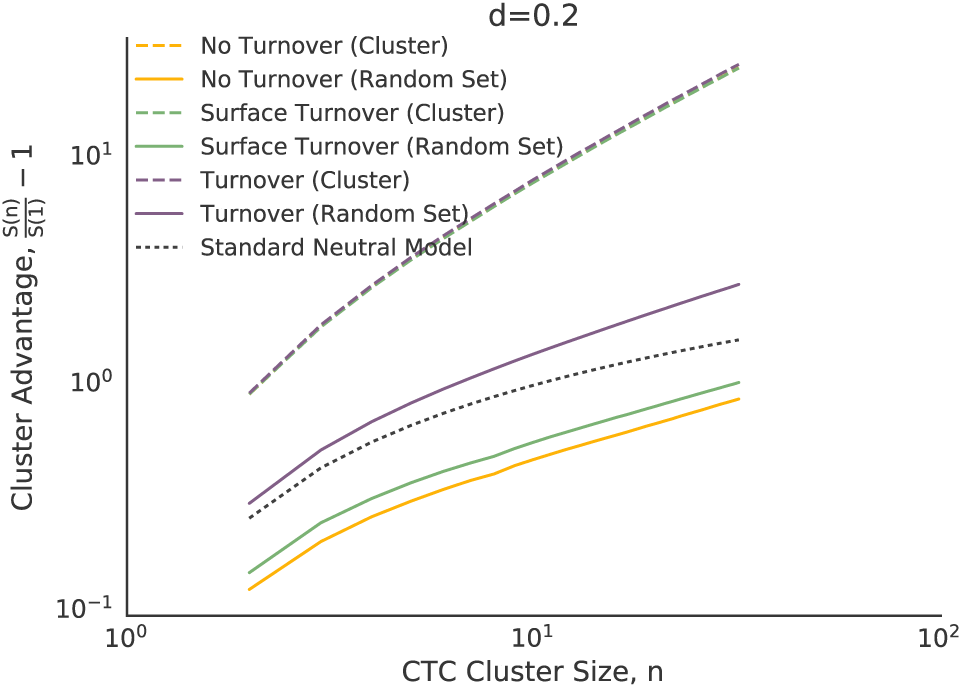
**Cluster advantage,** *A*(*n*)**, or the increase in number of distinct somatic mutations in a CTC cluster relative to single CTC**, as a function of cluster size for a random subset of 500 clusters drawn uniformly across the tumor. A law of diminishing returns applies to all models because of redundancy of mutations. The turnover model shows a 2-fold increase in the cluster advantage over the no turnover model. See also Fig S12 for *d ≤* 0.1.

**Figure 5:**
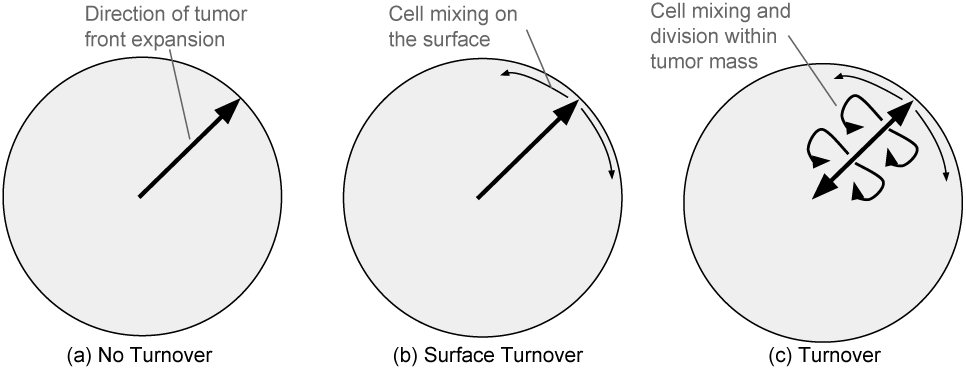
Serial founder effects and turnover explain spatial patterns of diversity. (a) In the no turnover model, the tumor front expands radially increasing genetic drift. There is little to no mixing and no divisions in the core: The number of somatic mutations increases with distance from the tumor center. (b) In the surface turnover model, the cells dying on the surface permit a small amount of mixing. This accounts for the higher number of somatic mutations per cluster. We still find increased diversity at the edge of the tumor because of the quiescent core. (c) In the turnover model, cells that die within the tumor can be replaced by cells from nighboring clones, leading to increased mixing and a supply of new mutations.

Whereas randomly sampled sets of cells show similar and almost linear increase of the cluster advantage with sample size, cell clusters show more variability. Turnover models have the highest cluster advantage, followed by the surface turnover model, and the no turnover model (Fig 4). Higher turnover increases the cluster advantage (Fig S12). Even low turnover with a death rate of *d* = 0.05 doubles the cluster advantage compared to the no turnover and surface turnover model (Fig S12).

## Discussion

### Global diversity

Even though tumor-wide distribution of allele frequencies in our simulations are consistent with Waclaw *et al.* (Waclaw et al. 2015), we reach opposite conclusions about the effect of cell turnover on genetic diversity. Waclaw *et al*. argued that turnover reduces diversity based on the observation that more high-frequency variants were observed in the tumor with turnover: A small number of clones make up a larger proportion of the tumor. Even though we can reproduce the observation, we find that turnover models in fact vastly *increase* diversity according to more conventional metrics, for example by increasing the number of somatic mutations (by ≈ 6.2× for *d* = 0.65) across the frequency spectrum. Both the increase in the number of dominant clonal mutations and the increased overall number of polymorphic sites have the same simple origin: A tumor model with turnover requires more cell divisions to reach a given size. Even though an early driver mutation has more time to realize a selective advantage and occupy a higher fraction of the tumor, carrier cells are also more likely to accumulate new mutations along the way leading to increased polymorphism (Fig 1 and Table S1). In other words, the Waclaw *et al.* metric of diversity (i.e., the number of clones above 10% in frequency) can reflect a higher concentration of common clones, but it is also confounded by changes in the mutation rate or in the number of cell divisions (i.e., an increase in the neutral mutation rate would counterintuitively result in a reduced measure of diversity).

At low rates of turnover, the global distribution of allele frequencies above 10^*−*4^ is well described by the Fusco *et al.* model assuming neutrality without turnover. With low turnover, the tumor is almost completely occupied, weakening the effect of selection (Fig S2): favorable mutations trapped within the tumor are hindered by spatial constraints (Fusco et al. 2016; Enriquez-Navas et al. 2016), whereas the effect of selection along the tumor edge is limited by the excess drift at the frontier (Excoffier, Foll, and Petit 2009). However, when turnover is increased to *d* = 0.65, the tumor is largely unoccupied (Fig S6) allowing for the release of the growth potential in fitter clones in the core.

### Spatial patterns in small clusters

The impact of turnover on cellular heterogeneity is more pronounced when considering small cell clusters (Figs 2 and S5). These fine-scale patterns can be interpreted by considering the expansion dynamics of each model and their impact on cell division and clonal mixing.

In all turnover models, the number of somatic mutations in a given cell is ≈ 3.0× higher at the edges than at the centre of the tumor, reflecting the higher number of divisions to reach the edge: The centre of the tumor is occupied early, which slows down cell division. Cells keep dividing due to turnover, however: For example, cells at the centre of the tumor with *d* = 0.2 have ≈ 8.4 somatic mutations, compared to ≈ 5.8 for the no turnover model. Turnover thus reduces, as expected, differences between edge and core cells: Without turnover, the number of somatic mutations per cell is ≈ 4.2 times higher at the edge than in the core, and the ratio is reduced to ≈ 2.0 when *d* = 0.2.

In the no turnover and surface turnover models, cell clusters show the same overall pattern of additional diversity at tumor edge. In the turnover model, however, we observe the opposite pattern: Even though edge *cells* still carry the most mutations, core *clusters* are now much more diverse than edge clusters. This can be understood in terms of a competition between the number of cell divisions (higher at the edge) and the effective population size (higher in the center). Even weak turnover vastly increases effective population size in the core. Even though a full analytical treatment of the spatial distribution of diversity in small clusters is beyond the scope of this article, the excellent agreement of the Fusco et al model predictions to global diversity patterns suggest that it provides an excellent starting point to build such a model. Supplementary Sections S.7, S.8, and S.9 provides simple order-of-magnitude estimate for the effects of clone mixing and late mutations (i.e., mutations in the core) observed on diversity patterns, including an extension of the Fusco et al model that accounts for late mutations. In addition to turnover rate, a key parameter that controls mixing is the expected distance between mother and daughter cells. In TumorSimulator, cells are allowed to reproduce to neighboring positions according to a Moore neighborhood, which leads to relatively diffuse small clusters. A challenge in building models for fine-scale diversity will be to implement realistic models of cell-cell interactions.

### Metastatic potential

The difference in somatic diversity between single CTCs and CTC clusters, measured through the cluster advantage, follows the expected law of diminishing returns: The more cells in the cluster, the fewer the number of unique mutations per cell. However, the trends vary by growth model and cluster origin. Cell mixing and rare, late mutations caused by turnover reduces neighboring cell similarity and increases cluster advantage.

Under the assumption that the presence or absence of a metastatic progression allele modulates metastatic potential of tumor cell clusters, the proportion of metastatic lesions that derive from circulating tumor cell clusters is highest in the turnover model. We can think of this as interference occurring between cells within a cluster. Alternately, this is an illustration of the advantage of not putting all one’s egg in the same basket, applied to tumor metastasis: Assuming that there is a chance component to cluster implantation, mixing due to turnover increases the likelihood that at least one virulence cell makes it to a hospitable site. Such an effect should be robust to details of the growth model.

In experiments, CTC clusters derived from primary breast and prostate tumors produced more aggressive metastatic tumors (Aceto, Bardia, et al. 2014) compared to single CTCs. This is likely due to differences in mechanical properties of the cluster or the creation of a locally favorable environment by the cluster, rather than by genetic differences. However, the present analysis suggests that this advantage can be enhanced by diversity within the cluster.

### Statistical power

Both fine-scale mixtures of cell phenotypes and clonally constrained mutations have been observed experimentally in tumors (Navin et al. 2010; Yates et al. 2015). Similarly, multi-region sequencing revealed high tumor heterogeneity in clear cell renal carcinoma (ccRCC) (Gerlinger, Horswell, et al. 2014) and esophageal squamous cell carcinoma (Hao et al. 2016), but low levels in lung adenocarcinomas (J. Zhang et al. 2014). This strongly suggests that the amount of mixing and late mutations varies substantially across tumors, with ccRCC data being better described by a model with turnover, whereas lung adenocarcinoma data more closely resembles a model with low or no turnover.

In practice, distinguishing between mixing, turnover, mutations, and tumor growth idiosyncrasies will be challenging. Among limitations of our model, we note the assumption of spherical tumor shape and the absence of complex physical contraints (which HCC tumors may experience). Another limitation of the present model is the rigid computational grid which prevents cells from pushing each other out of the way, which constrains growth in the center of the tumor. This constraint plays a role in reducing diversity at the center of the tumor, but it may not be realistic in the earlier stages of tumor growth. The importance of such effects is largely unknown, and it is likely to vary between tumors and tumor types. Fortunately, we have shown that we are at the cusp of being able to test such models quantitatively. A sampling experiment with twice as many samples than were collected in the HCC patient studied above would enable us to either validate or reject the current state-of-the-art models confidently (Fig 3b). Alternatively, sequencing of small clusters would further allow us to discriminate between the different models of turnover.

In either case, the use of frequency cutoffs can strongly affect inferred spatial patterns of diversity: a focus on common variants means a focus on old branches of the tree (Fig S10). This emphasizes the mean number of divisions per cell, which is larger at the edge, but fails to capture recent mutations and clonal mixing, which have larger impact at the core. Thus spatial patterns inferred using variants at frequency above 1% are more similar across models, and can be opposite to those including all mutations (Fig S11).

Data collection schemes including the lung TRACERx study (Jamal-Hanjani, Hackshaw, et al. 2014; Jamal-Hanjani, Wilson, et al. 2017) will help us put the state-of-the-art models to the test and identify such important parameters of tumor growth. Given our power analysis, we find that sequencing small contiguous cell clusters provides a richer picture of tumor dynamics compared to larger biopsies, with little to no loss in power, assuming that few-cell sequencing can be performed accurately.

## Conclusion

This work set out to answer two simple questions: First, should we expect substantial heterogeneity at the cellular scale within tumors and within circulating tumor cell clusters? The answer to the first question is most likely yes, as even the models with no turnover exhibit measurable cluster heterogeneity.

The second question was whether this heterogeneity, sampled through liquid biopsies or multi-region sequencing, is informative about tumor dynamics. Given that state-of-the-art models produce very different predictions about the level of cluster heterogeneity, the answer is also positive. This work identified some of the key factors that determine cluster diversity, especially the interaction between range expansion and cell turnover leading to late mutations and mixing. Even if no diversity were observed at all in CTC clusters, it would enable us to reject the present models in favor of models including additional biological factors that favor the clustering of genetically similar cells. Measuring diversity, or the lack of diversity, within circulating tumor cell clusters or fine-scale multi-region sequencing is therefore a promising tool for both fundamental and medical oncology.

## Author Contributions

Conceptualization, S.G.; Methodology, S.G.; Software, Z.A.; Investigation, Z.A. and S.G.; Writing Original Draft, Z.A.; Data Curation Z.A.; Review & Editing, Z.A & S.G.; Visualization, Z.A.; Funding Acquisition, Z.A. and S.G.; Resources, S.G.; Supervision, S.G.

## Acknowledgments

We thank Julien Jouganous, Hamid Nikbakht, Yasser Riazalhosseini, Aaron Ragsdale and Robert Sladek for useful discussions. This research was made possible thanks to a Canadian Institutes of Health Undergraduate Research Award in computational biology, funding reference numbers 139962 and 145987 and Frederick Banting and Charles Best Canada Graduate Scholarship. This research was undertaken, in part, thanks to funding from the Canada Research Chairs program and a Sloan research fellowship.

## S Supplemental Information

### S. 1 Tumor growth model

The tumor consists of cells that occupy points on a 3D lattice. Empty lattice sites are assumed to contain normal cells which are not modelled explicitly in TumorSimulator.

Each cell has an associated list of genetic alterations which represent single nucleotide polymorphisms (SNPs) that can be either passenger or driver. Driver mutations increase the growth rate by a factor 1 + *s*, where *s ≥*0 is the selective advantage of a driver mutation.

At *t* = 0, the simulation begins with a single cell that already has an unlimited growth potential.

The TumorSimulator algorithm then proceeds to grow the tumor through the following steps:

1. Select a random cell to be the mother cell.
2. Set the cell birth rate to *b^′^* = *b*_0_(1 + *s*)^*k−k*^_max_, where *b*_0_ is the initial tumor birth rate, *s* is the average selective advantage of a driver mutation, *k* is the number of driver mutations present in the mother cell and *k*_max_ is the maximum number of drivers in any cell.
3. Randomly select a lattice point adjacent to the mother cell. If empty, create a genetically identical daughter cell at that position with a probability *b^'^*. If no cell created, or no empty sites are found proceed to 5.
4. Independently give mother and daughter cells additional passenger and driver mutations. The number of passenger and driver mutations are drawn according to Poisson distributions with mean *µ_p_* and *µ_d_,* respectively, and are drawn independently for the mother and daughter cell. Each mutation is unique and there is no back-mutations or recurrent mutations.
5. Kill (i.e., remove) the mother cell with probability *d*(1 + *s*)^*−k*^_max_.

In our analysis, we consider three turnover scenarios corresponding to three values of the death rate *d*: (i) No turnover (*d* = 0), corresponding to simple clonal growth (Hallatschek et al. 2007); (ii) Surface Turnover (*d*(*x, y, z*) *>* 0 only if *x, y, z* is on the surface), corresponding to a quiescent core model (Shweiki et al. 1995) (iii) Turnover (*d >* 0 everywhere), a model favored in Waclaw et al. 2015 to explore ITH.

The initial birth rate (*b*_0_ = ln(2)), driver mutation rate *µ_d_* = 2 × 10^*−*5^, and selective advantage (*s* = 1%) were kept consistent with Waclaw et al. 2015 except where otherwise noted. In addition to varying the turnover model (full, surface, or none), we vary its intensity by controlling the death rate, *d ∈* {0.05, 0.1, 0.2, 0.65}. TumorSimulator also has a parameter that controls migration of cells to form new independent cancer lesions. We did not allow such local migrations, as they would have little effect on the very fine-scale diversity in the primary tumor. We used two values for the passenger mutation rate: *µ_p_* = 0.01 to facilitate comparison with simulations from Waclaw et al. 2015 (Waclaw *et al.* simulated with *µ_p_* = 0.01, but reported a mutation rate of 0.02 to account for an equivalent rate per diploid genome), and *µ_p_* = 0.01875 to match experimental observations from Ling et al. 2015 (Since the number of passenger mutations grows linearly with the mutation rate, we simply scaled *µ_p_* based on the difference between predictions using *µ_p_* = 0.01 and the data from Fig 3a.) All tumors were grown until they had 10^8^ cells except where otherwise stated.

TumorSimulator (Waclaw et al. 2015) is available at http://www2.ph.ed.ac.uk/bwaclaw/cancercode/.

### S. 2 CTC cluster synthesis

Experimental evidence suggests that CTC clusters are formed from neighboring cells in the primary tumor and not by agglomeration or proliferation of single CTCs in the blood (J. M. Hou et al. 2012; Aceto, Bardia, et al. 2014). To represent circulating tumor cell clusters, we therefore sampled spherical clusters (with a large radius) of cells in different areas of the tumor produced by the Waclaw *et al.* model. To get a fixed number of cells in the cluster, *n*, we picked the *n* closest cells to the center-of-mass of this sphere. We varied the number of cells in the cluster from *n* = 2 to *n* = 30 to represent the range of empirical findings (Marrinucci et al. 2012).

### S. 3 Power Analysis

To establish the effectiveness of sequencing CTC clusters versus larger biopsies at detecting a trend and distinguishing between models, we conduct a power analysis. We use linear regression on the number of somatic mutations per cluster (or biopsy) of size *n* as a function of distance *r* from the tumor center-of-mass (i.e, *S*(*n, r*) = *mr* + *c* where *m* and *c* are regression coefficients). Clusters and biopsies to regress are sampled at random from a previously generated set of 1000 samples. Given a sample size and cluster size, we resample 100 subsets from these 1000 samples to estimate proportion of regressions that were significant (*p <* 0.01). To capture the direction of the slope, we calculate the sign of the coefficient *m* and report the *signed* proportion of significant regressions. For larger biopsies, we apply a frequency cutoff and only includes a mutation in the analysis if it is above a certain cluster-wide frequency, thus simulating the mutant allele frequency cutoff from sequencing experiments (Ling et al. 2015).

### S. 4 Standard Neutral Model for Cluster Advantage

The relative increase in the number of distinct somatic mutations in a CTC cluster versus a single CTC is given by the *cluster advantage*, i.e., 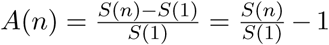, where *S*(*n*) is the number of somatic mutations in a cluster of size *n* and *S*(1) is the number of somatic mutations in the cell closest to the center-of-mass of the cluster (as described in Section CTC cluster synthesis). A higher cluster advantage indicates that a CTC cluster is more potent relative to a single CTC from the same tumor. In other words, a higher cluster advantage means less genetic redundancy within a cluster. Under the standard neutral model (infinite sites, neutral evolution, random mixing), and therefore the expected number of somatic mutations is 𝔼(*S*(*n*)) = *µH*(*n −* 1) (Durrett 2008), where *H*(*n*) is the *n*-th harmonic number, 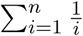.

### S. 5 A geometric model

To estimate the frequency distribution of common variants, we model the tumor as a continuously growing sphere where only surface cells divide. If a mutation appears in a cell at the surface of the tumor at a time when the tumor has radius *r*, we suppose that this mutation occupies a crosssection area *a*^2^ of the tumor surface. It therefore occupies a fraction 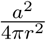 of the surface of the tumor at that point. If the tumor grows radially outwards and reaches a radius of *R*, the descendants of this cell occupy a fraction 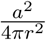 of the space yet to be occupied, and the mutation itself will occupy a fraction

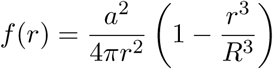

of the final tumor, which is the volume of a spherical cone with its tip removed. We can then integrate over all possible radii *r* where mutations occur. The density *ρ*(*r*) of mutations occurring at radius *r* is proportional to the density of cells at that locus

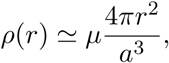

with *µ* the mutation rate per cell. The frequency spectrum is therefore

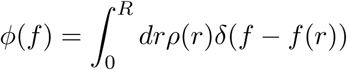

If we focus on common mutations, which occurred at *r ≪ R*, we can approximate 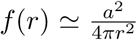, leading to

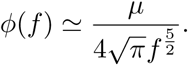

We show in the next section that a model accounting for stochastic fluctuations in the early reproductive success of a mutation, or weak changes in selection, preserves this scaling behavior, but with an overall scale factor *ζ* that depends on details of the growth model, i.e.

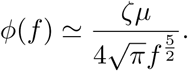

Fig 1 shows the agreement of simulation results to the geometric model with *ζ* = 30 for high frequency mutants. As mentioned above, variants at less than 1% frequency follow a distinct power law with slope closer to our estimate of 1.61, which is similar to the theoretical value of 1.55 described in Fusco *et al.* (2016).

### S. 6 Allele frequency distribution under a stochastic spherical growth model

The deterministic model presented above does not take into account the stochastic variation in the fate of cells, which is especially important in the first few generations after a mutation appears. To account for this, we can imagine that the initial frequency of each new mutation gets multiplied by a random factor *i* to account for the random differences in success in the original cells over the first few generations. In other words, *i* is the number of descendants produced by the original cell divided by the expected number of descendants for other cells at the same radius. If we only consider mutations with given *i*, we find

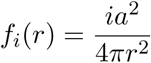

and

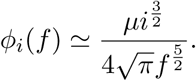

If we assume that multipliers are drawn from a probability distribution *P* (*i*) that is independent of *r*, we get an expected frequency spectrum

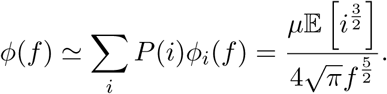

Even though the 5*/*2 scaling behavior is maintained, the expectation 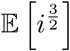 can be much larger than 1, as there is an early settler advantage in this model. However, the value of this scaling factor depends on the details of the growth model (Fig 1 and S2).

More generally, the 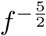 asymptotic result is derived under an extremely simple model. The Fusco et al. model (Fusco et al. 2016) captures a very similar scaling, but with a much more detailed model of stochastic fluctuations that captures both rare and common variant scaling. Neither models take into account selection and turnover. Analytical results under selection are difficult to obtain because moment-based approaches that close under neutrality do not close under selection (see, e.g.,Weinstein et al. 2017; Korolev et al. 2010; Jouganous et al. 2017).

### S. 7 Expected frequency of a mutation in a given cell

Following the Fusco *et al* model, the distribution of allele frequencies can be approximated by its asymptotic values

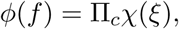

where 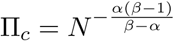, 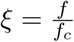 and 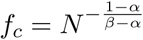 is the transition point between the two asymptotic regimes. Finally

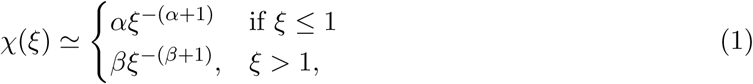

where *N* is the number of cells in the tumor and *α* = 0.55 and *β* = 2.3 are scaling factors that depend on the geometry of tumor growth.

Using this approximation, we can compute basic statistics for the expected frequency of sampled alleles. For example, the expected allele frequency of a mutation selected uniformly at random is

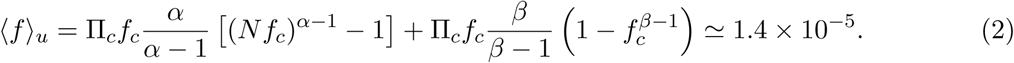

If we sample mutations proportionally to their population frequency, we get

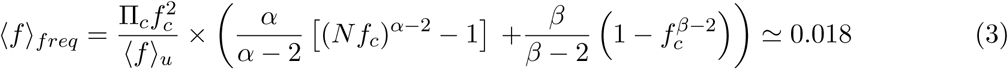

so that the expected frequency of a mutation observed in a given cell is at reasonably high frequency. That is to say that the typical clone size, in a tumor of size 10^8^, is approximately 1.8 × 10^6^.

Similarly, the the probability of drawing a mutation at frequency *f*, given that mutations are sampled according to their frequency, is

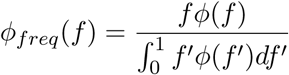

and the cumulative distribution function of the allele frequencies for mutation drawn proportionally to the allele frequency is

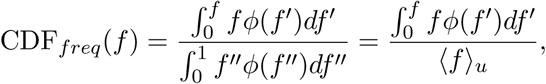

from which we infer that less than 2% of variants in a cell drawn at random are derived from clones with frequency below 10^*−*5^: Over 98% of cells derive from clones of size over 1000.

### S. 8 Number of cell divisions, and properties of the tumor core

We would like to estimate the average number of divisions since tumor beginning that cells at a given position in the tumor have undergone. We consider a two-stage model, wherein we first have straightforward tumor expansion which can be described by the Fusco et al ‘bubble and sector’ model, and subsequent alteration of this state under a steady-state model. In the first stage, the main effect of turnover is to increase the number of divisions necessary for the tumor necessary to reach a given radius *R*. Under turnover, it takes more divisions for the tumor to reach a given radius, and we find empirically that the number of divisions required to reach a given radius increases approximately by a factor (1 + *d*) for low turnover.

We’ll distinguish between ‘early’ and ‘late’ mutations according to whether a mutation occurred on the expansion front (early), or behind the front (late). To estimate the rate of division within the tumor core, we must first estimate the unoccupied cell density *e* within the tumor. This can be estimated as close to 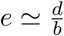 by assuming that growth due to births *eb* is offset by death *d*. (It is approximate because it assumes that the probability of drawing an empty cell next to the selected mother cell is *e* — this is not exact if there are spatial correlations in cell occupancies.)

The final radius of the tumor is therefore approximately 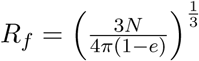. This is close enough to the observed values in Fig 2 (for example, this predicts *R_f_* = 288 for *d* = 0 and *R_f_* = 323 for *d* = 0.2).

To estimate the number of late mutations, we first need to compute the expected number of divisions occurring along a given lineage after the tumor front has passed. This can be estimated by first considering the expected number of times a given core cell is selected while the tumor grows from radius *R* to *R* + 1. In a model where the tumor has a smooth boundary, a cell on the boundary has probability *γb* of reproducing successfully (i.e., we have a probability *γ ≃* 1*/*2 of drawing an empty site nearby, and probability *b* of successfully reproducing on that site).

Now, while growing from size *R* to size *R* + 1, we consider that each cell on a surface of area 4*πR* must successfully reproduce on average 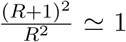 times. It must therefore be selected 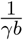 times, on average, for the tumor to move forth one unit. Since cells are chosen at random, each cell inside the tumor must be picked, on average, the same number of times as edge cells. This leads to, on average, 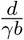 deaths (and, at equilibrium, the same number of births).

Thus the total number of births/deaths per occupied cell at distance *R*_0_ after the front has passed is

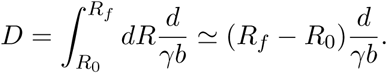

For *d* = 0.2, and *R_f_* = 323 this means approximately 188 deaths and birth per occupied cell. The expected number of mutations on a lineage increases by *µ ≃ ×*10^*−*2^ with each birth/death cycle. Thus each lineage gains order of two new mutations in the core. This is consistent with the lineages drawn on Fig S10, and with the increase in the number of clones per cell in Fig 2.

Clones derived from these mutations are extremely unlikely to reach frequencies comparable to the bubble and sector clones contributing to diversity in the Fusco et al. model. Thus weak turnover therefore induces a third, distinct regime of late clones, in addition to the bubbles and the sectors, which will remain very rare. Late clones will have a higher relative impact near the center of the tumor, given the additional time for late clones to develop, and the reduced number of early clones.

Because the core is near birth-death equilibrium, the expected number of descendants of a given cell (and therefore of a new mutation) is one. Thus the expected number of late mutations in a cell at distance *R* of the core is simply 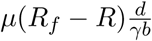.

### S. 9 Mean frequency of late clones and cluster diversity

If we suppose that late clones remain very small, we can model each cell division as independent of each other. That is, we can neglect the probability that mutant cells replace each other and model clone growth as a critical Galton-Watson branching process a probability *d* of dying or branching. This apparently coarse approximation is reasonable here because TumorSimulator uses a Moore neighborhood with 26 neighbors: the fact that a mother cell occupies one of these 26 neighborhood cells has a low impact on the probability of the daughter cell to divide. Further divisions will not crowd out space as long as the clusters remain relatively small: a cell’s daughter will be at approximate mean squared distance 3 * (18*/*26) = 2.1 (the factor of three accounts for the three dimensions, and 18*/*26 is the mean squared displacement in each direction. This is approximate because displacement along the three directions is not independent). The grand-daughter will be at mean squared distance 4.15, and so forth. A simple toy model where cells carrying a mutation are allowed to divide into neighboring grid points with probability *e*, irrespective of occupancy (i.e., grid points can carry multiple cells), shows relatively little overlap for the parameter ranges studied here: For a mutation occurring at the founding of the tumor with parameters *d* = 0.1, *e* = 0.14, and *R_f_* = 303, there is only 6% overlap on average by the time the tumor has size 10^8^ (i.e., the mean number of occupied gridpoints is only 6% lower than the number of cells, including those jointly occupying a grid-point.)

A very crude estimate of the number of segregating sites in small clusters can therefore be obtained by assuming that late clones are in fact so diffuse that is is unlikely that a small cluster will capture more than one clone cell – this will naturally overestimate *S*(*n*) for large clones and large clusters but we find that it is an appropriate approximation for small clusters, or for parts of the tumor that experienced relatively few late divisions (Fig S7). In Fig S7, predictions are obtained by using the empirical number of early mutations observed in single cells under no turnover (*S_d_*_=0*,n*=1_(*R*)), shown as a dotted line, scaling it by the empirical factor (1 + *d*) discussed in the previous section, and adding the predicted number of late mutations 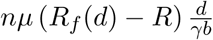.

We computed above 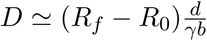, the estimated number of cell divisions per occupied site between the front passage and present. Mutations accumulate at a constant rate during this time, and so the typical late mutation at this position will only have 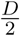 generations to experience genetic drift. For *d* = 0.2, and *R_f_* = 323, this means 94 cycles.

To model the distribution of clone size, we consider the Galton-Watson model with variance 2*d*(1 − d). The variance in clone size in the Galton-Watson model after *j* generations is simply 2*d*(1 − d)*j*.

We can estimate the distribution of surviving sizes using Yaglom’s asymptotic limit (Lyons, Pemantle, and Peres 1995), and find that the size distribution of surviving lineages after *j* generations is

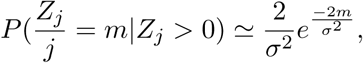

and

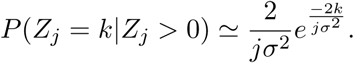

The expected size of a surviving clone is approximately

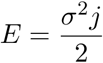

and the asymptotic survival probability is simply 1*/E,* per Kolmogorov’s estimate.

Thus the overall probability of having a clone of size *k >* 0 given *j* steps is

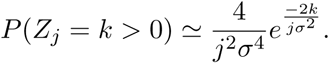

Finally, we must add contributions from all mutations appearing at all positions in the tumor. If we imagine that each cell in the tumor contributes mutations at constant rate *µ*, from the moment the front crosses it, then the number of mutations with an expected *j* death cycles is

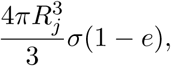

where *R_j_* is the maximal radius for which mutations can have an expectation of going through *j* death-birth cycles.

Thus we simply need to sum the number of late mutations occurring over all positions in the tumor. Through each cycle *R* to *R* + 1, there are 4*πR*^3^(1 − e)*/*3 cells in the tumor, and an average of 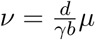 mutations that will appear, each of which will survive on average 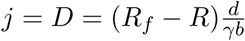 generations. Thus

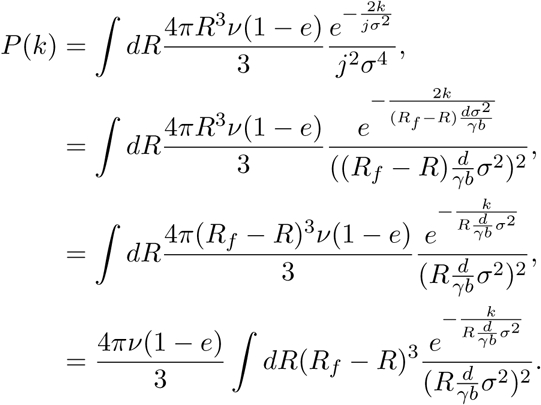

This can be integrated using Mathematica

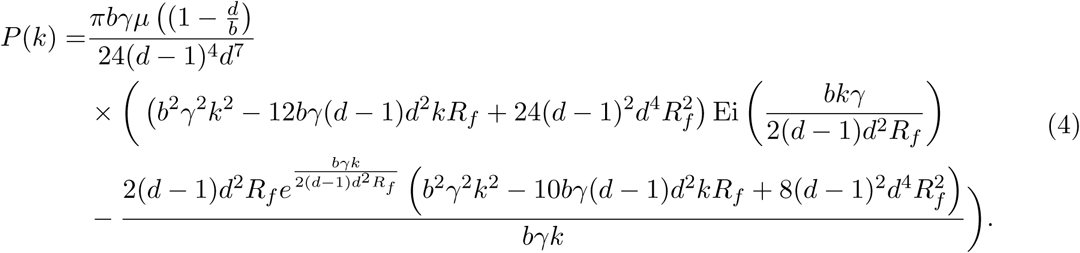

This provides a good estimate for the excess of rare variants observed in *d* = 0.1 and *d* = 0.2 compared to *d* (Fig S4b).

### S. 10 Code Availability

The code to reproduce simulations, analyses and figures can be found at https://github.com/zafarali/tumorheterogeneity. Parameters for each simulations and details of how to reproduce results and figures are specified in Table S2.

**Table 1:**
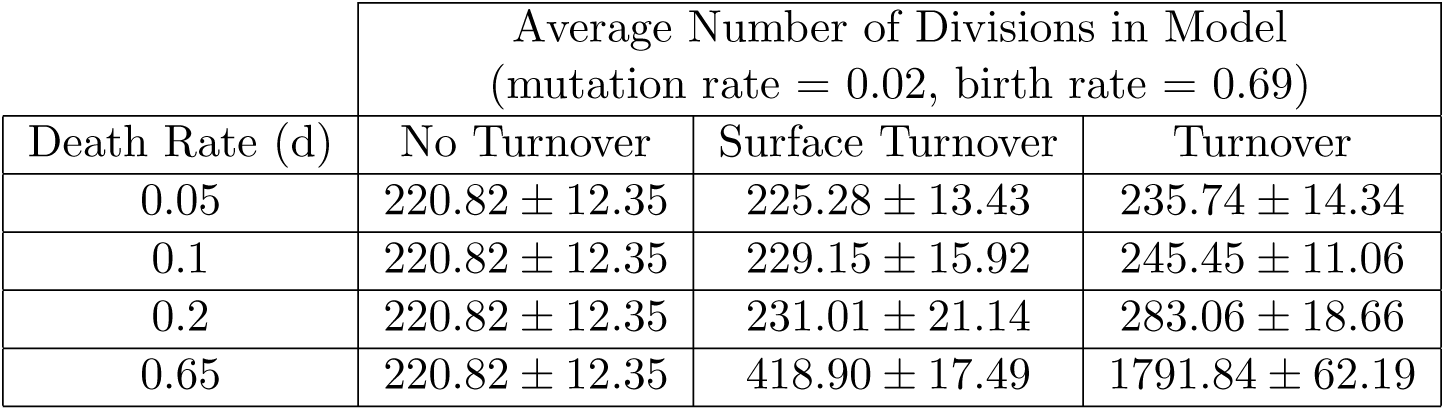
Average number of generations for a cell in each model (estimated from the number of somatic mutations per cell divided by the mutation rate, *µ* = 0.01). Standard deviation in brackets. The number of divisions increases with the death rate.

**Supplemental Figure 1:**
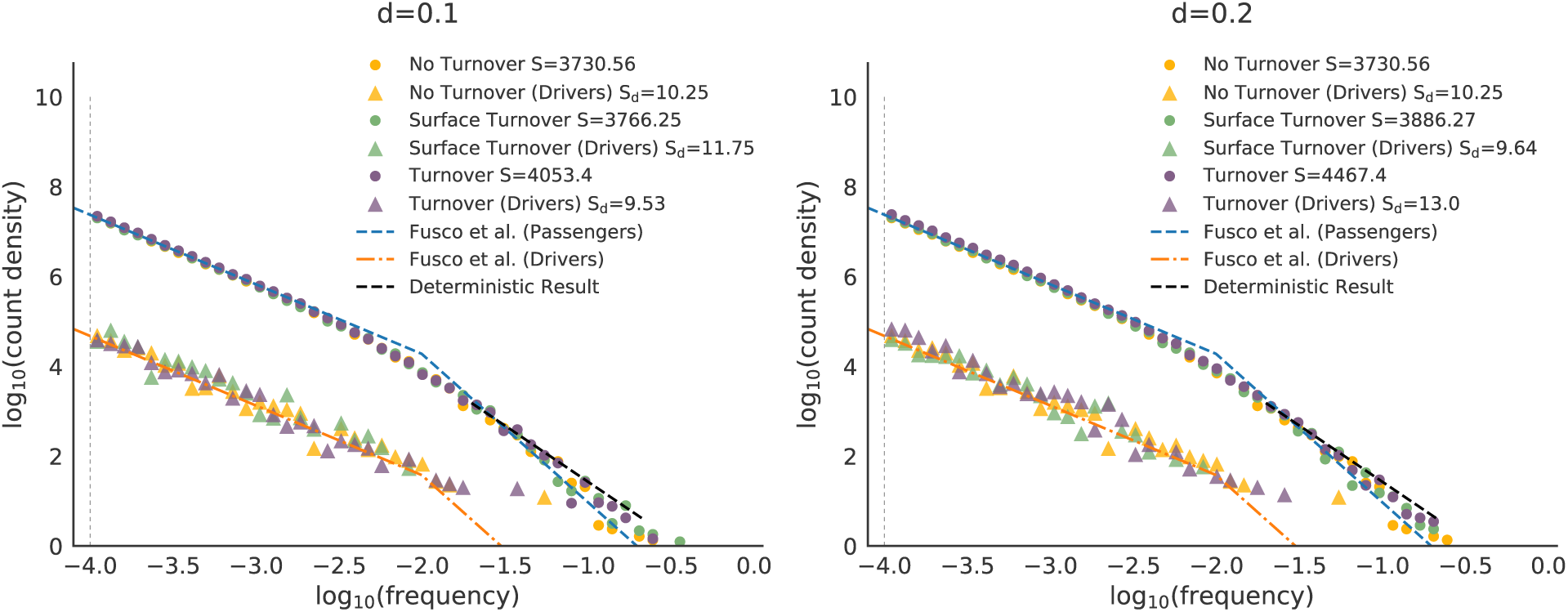
Allele frequency spectra for low death rates, *d ∈* {0.1, 0.2} show similar scaling laws. Total allele frequency distribution is shown using circles and driver frequency distribution using triangle. The total number of somatic mutations, S, and the total number of driver mutations, *S_d_*, in the tumor is shown in the legend (average of 15 simulations). The vertical gray dotted line shows the minimum frequency of mutations returned by TumorSimulator. The black dotted line shows the asymptotic result of a geometric model with a scaling of *ζ* = 30 and is described in Supplementary Section S.5. The blue and oranged dashed lines shows the result from Fusco *et al.*. See Fig 1 for *d* = 0.05 and *d* = 0.65.

**Supplemental Figure 2:**
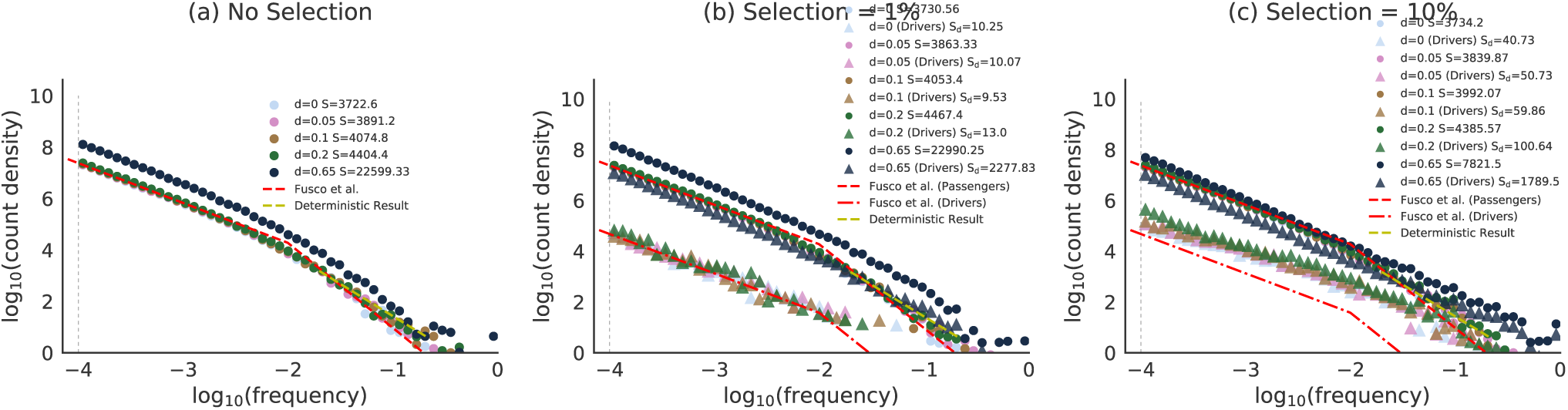
Comparison of the allele frequency spectra for simulations with selection rates (a) *s* = 0, (b) *s* = 1% and (c) *s* = 10% for different death rates *d*. The allele frequency spectra are similar across selection coefficients at *d ∈* {0, 0.5, 0.1, 0.2}. Only under high turnover (*d* = 0.65) is there a departure from the no turnover scaling result of Fusco *et al*.

**Supplemental Figure 3:**
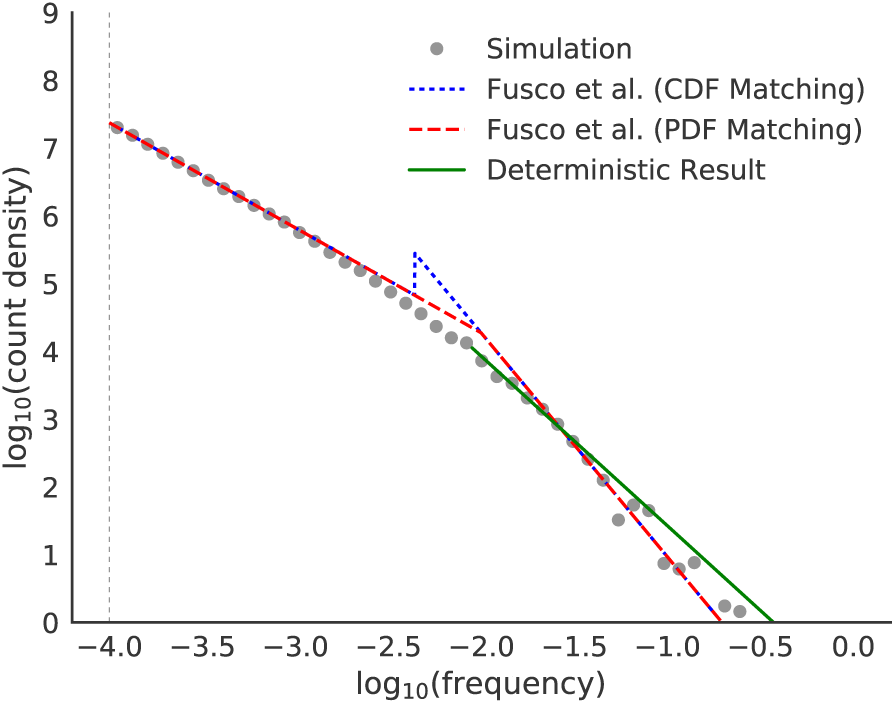
Continuity matching in probability space. Blue dotted line represents the solution from Fusco *et al.* with two scaling regimes described by the powers *α* = 0.55 and *β* = 2.3. Fusco *et al.* imposed continuity matching on the cumulative distributions of frequencies (CDF), leading to *f*_*c*_ = 10^−2.06^. The resulting probability distribution of afrequencies (PDF) is the derivative of the cumulative function and thus discontinuous. 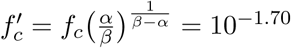, where α = 0:55 and β = 2:3 are the low and high frequency Continuity matching in frequency scaling factors respectively. Gray circles show results from a simulation with a neutral selection coefficient and no turnover. The green solid line shows the deterministic geometric model with a scaling of *ζ* = 30.

**Supplemental Figure 4:**
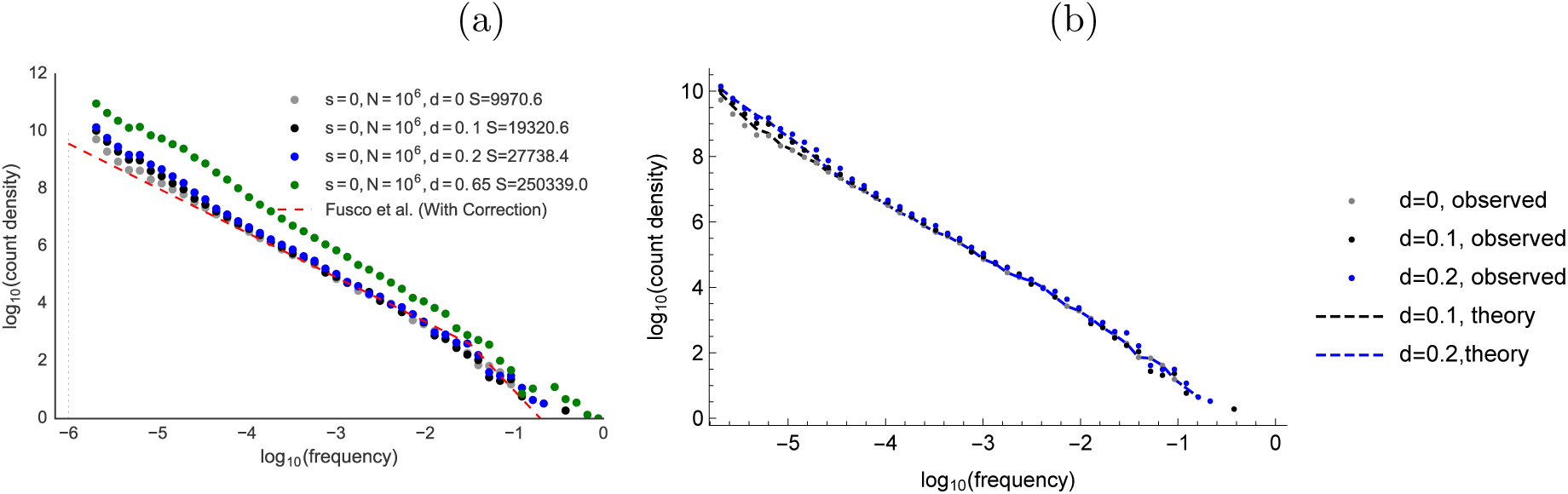
(a) Effect of turnover on rare variant frequency distribution, showing departure from the (no turnover) Fusco et. al analytical model. We simulate a smaller tumor with *N* = 10^6^ to make it computationally tractable to list all mutations in the tumor. (b) Validation of the theoretical model from Section S.9: the excess of rare variants for *d* = 0.1 and *d* = 0.2 can be estimated using a Galton-Watson model of clonal growth. The dashed lines are obtained by adding the prediction for the distribution of late clones from Eq 4 to the observation with *d* = 0.

**Supplemental Figure 5:**
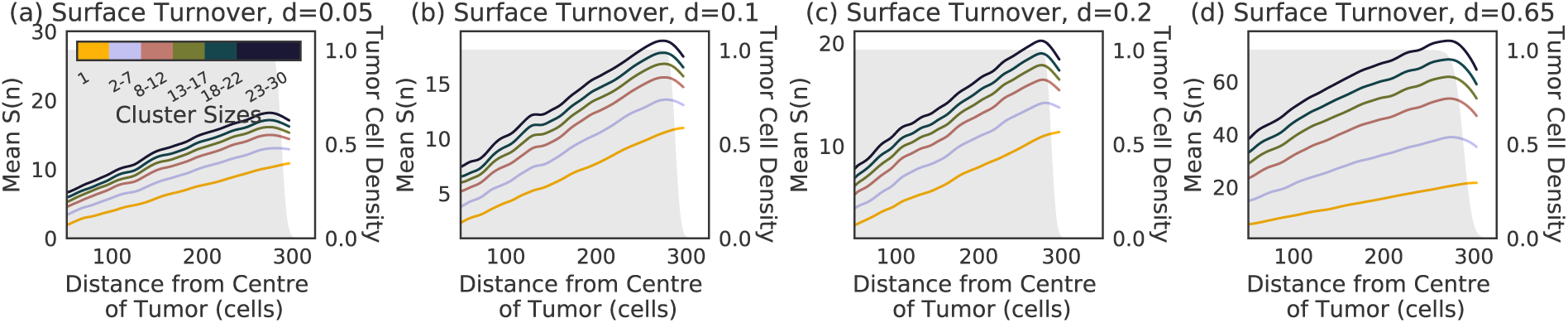
Spatial distribution of the number of somatic mutations per cluster in the surface turnover model with death rates. **(a)** *d* = 0.05, **(b)** *d* = 0.1**, (c)** *d* = 0.2 **and (d)** *d* = 0.65. Trends are similar to the no turnover model indicating that a majority of the effects seen in the turnover models is due to the fact that cell death and mixing can occur throughout the tumor. See Fig 2a for *d* = 0 and Fig 2b-d for the corresponding turnover models.

**Supplemental Figure 6:**
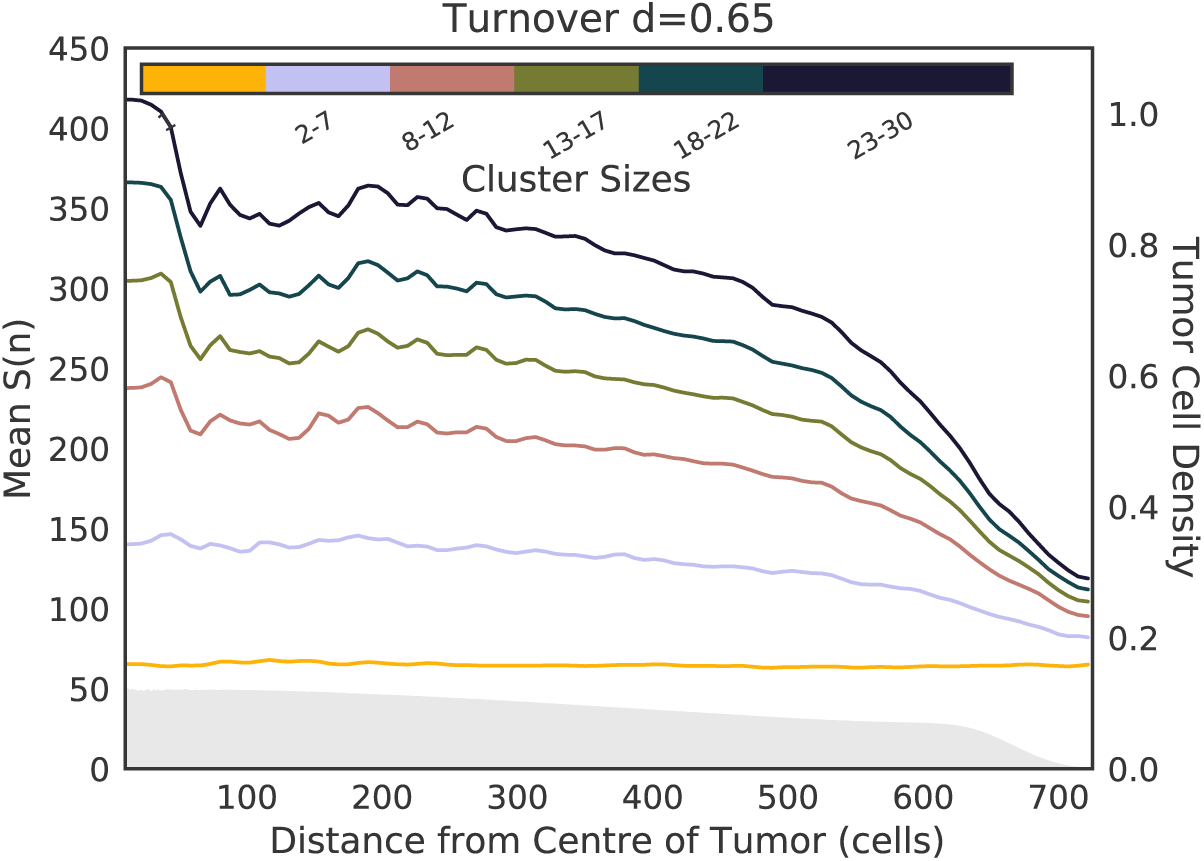
**Spatial distribution of the number of somatic mutation per cluster in a turnover model with** *d* = 0.65. Large clusters show a stronger decreasing *S*(*n*) with distance from the centre of the tumor compared to lower death rates (Fig 2). See Fig 2 for simulations with *d <* 0.65 and Fig 5 for the surface turnover model.

**Supplemental Figure 7:**
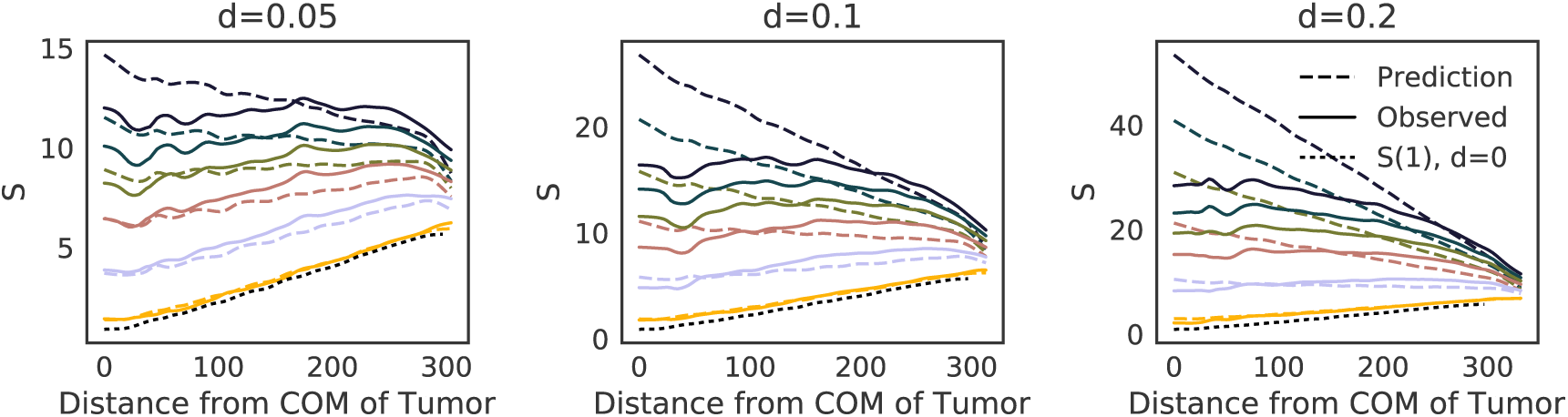
Order-of-magnitude estimates from Supplementary Section S.9 for the number of somatic mutations per cluster for different turnover models and their agreement with simulations. Colors are consistent with Fig 2, S5 and S6

**Supplemental Figure 8:**
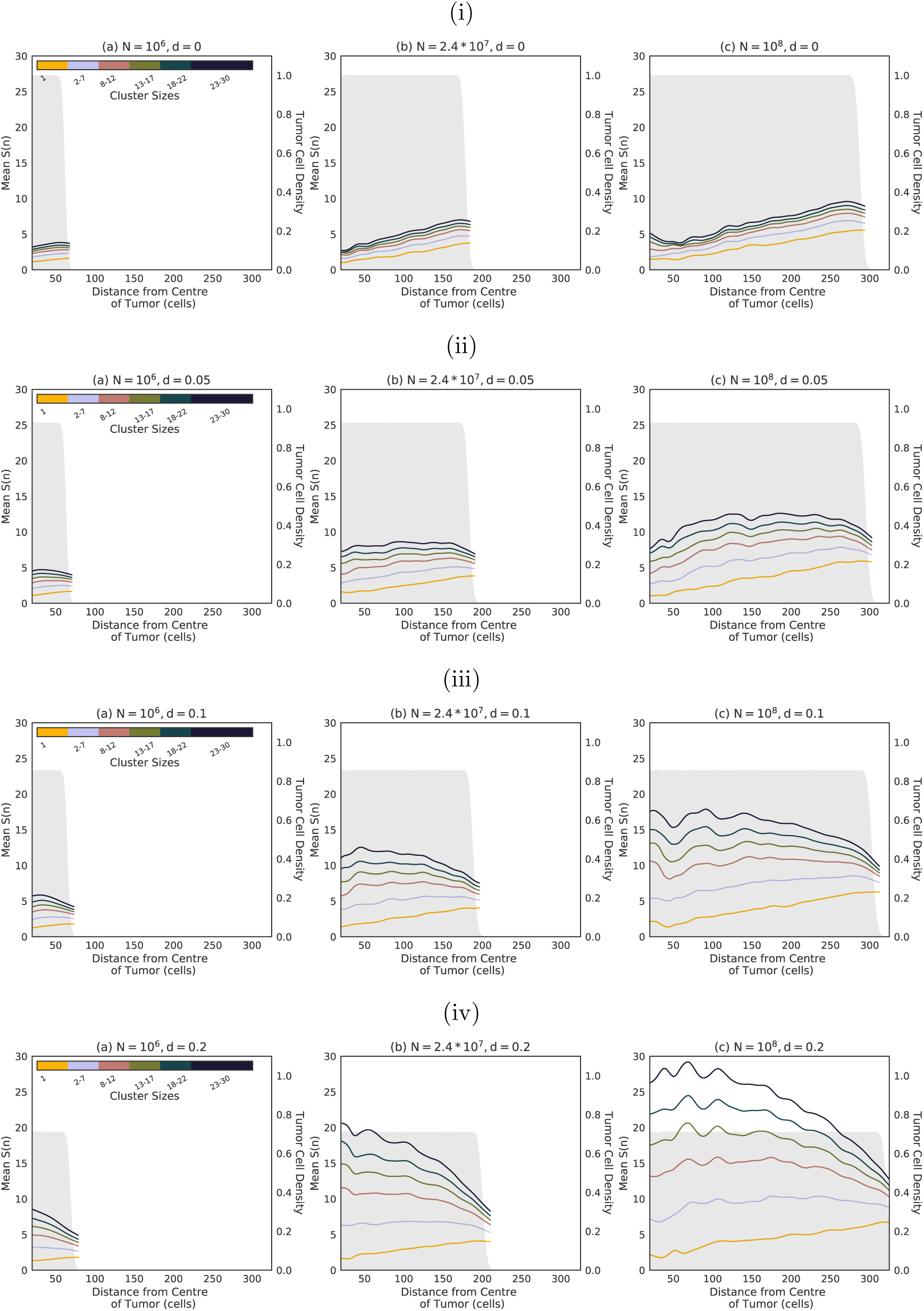
Time course view of the spatial distribution of the number of somatic mutations per cluster as the tumor grows from 10^6^ to 2.4 × 10^7^ and 10^8^ cells for (i) *d* = 0, (ii) *d* = 0.05, (iii) *d* = 0.1 and (iv) *d* = 0.2.

**Supplemental Figure 9:**
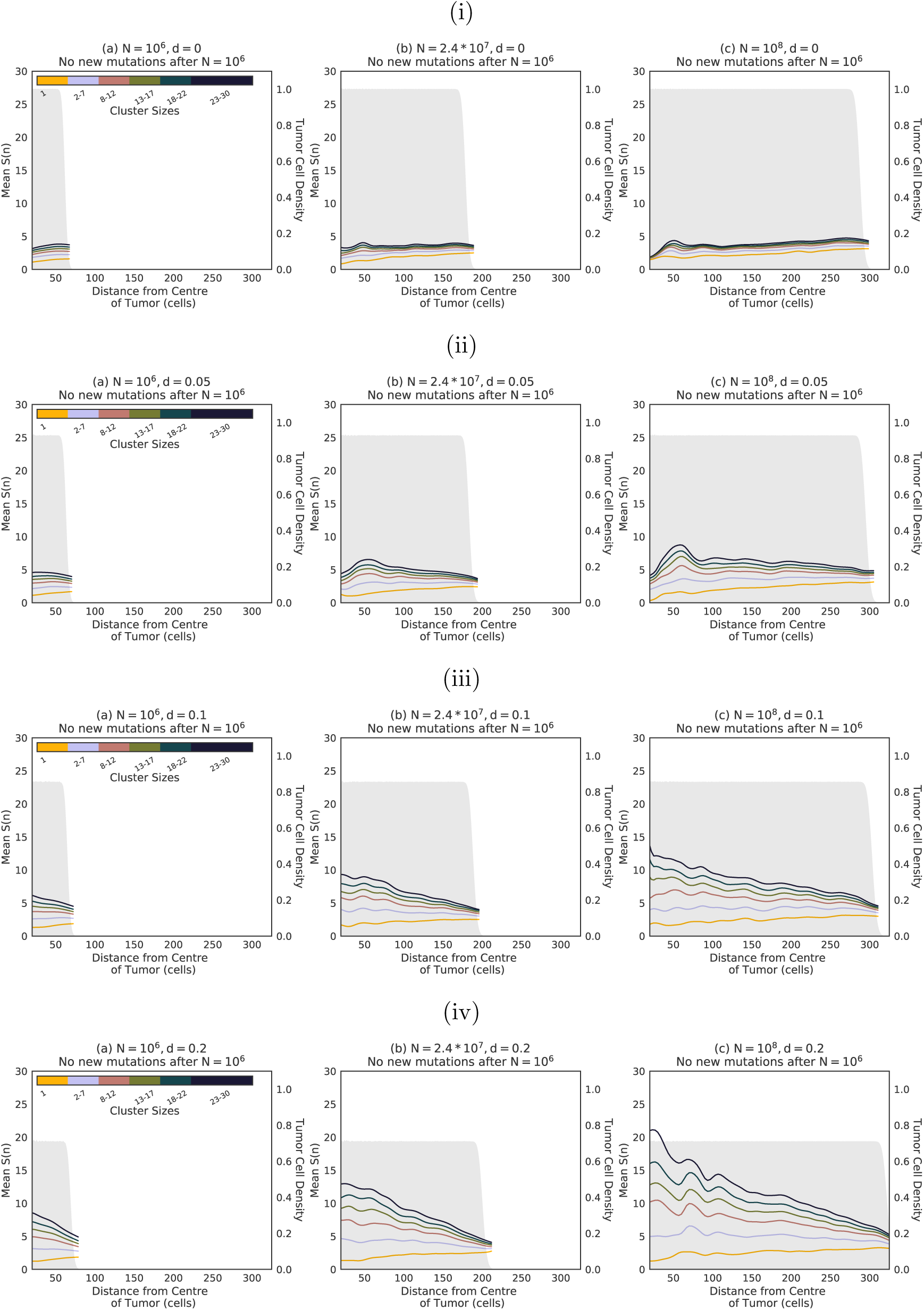
Time course view of the spatial distribution of the number of somatic mutations per cluster as the tumor grows from 10^6^ to 2.4 × 10^7^ and 10^8^ cells for for (i) *d* = 0, (ii) *d* = 0.05, (iii) *d* = 0.1 and (iv) *d* = 0.2 if no new mutations are created when the tumor reaches 10^6^ cells, thus revealing the contributions of clonal mixing and genetic drift.

**Supplemental Figure 10:**
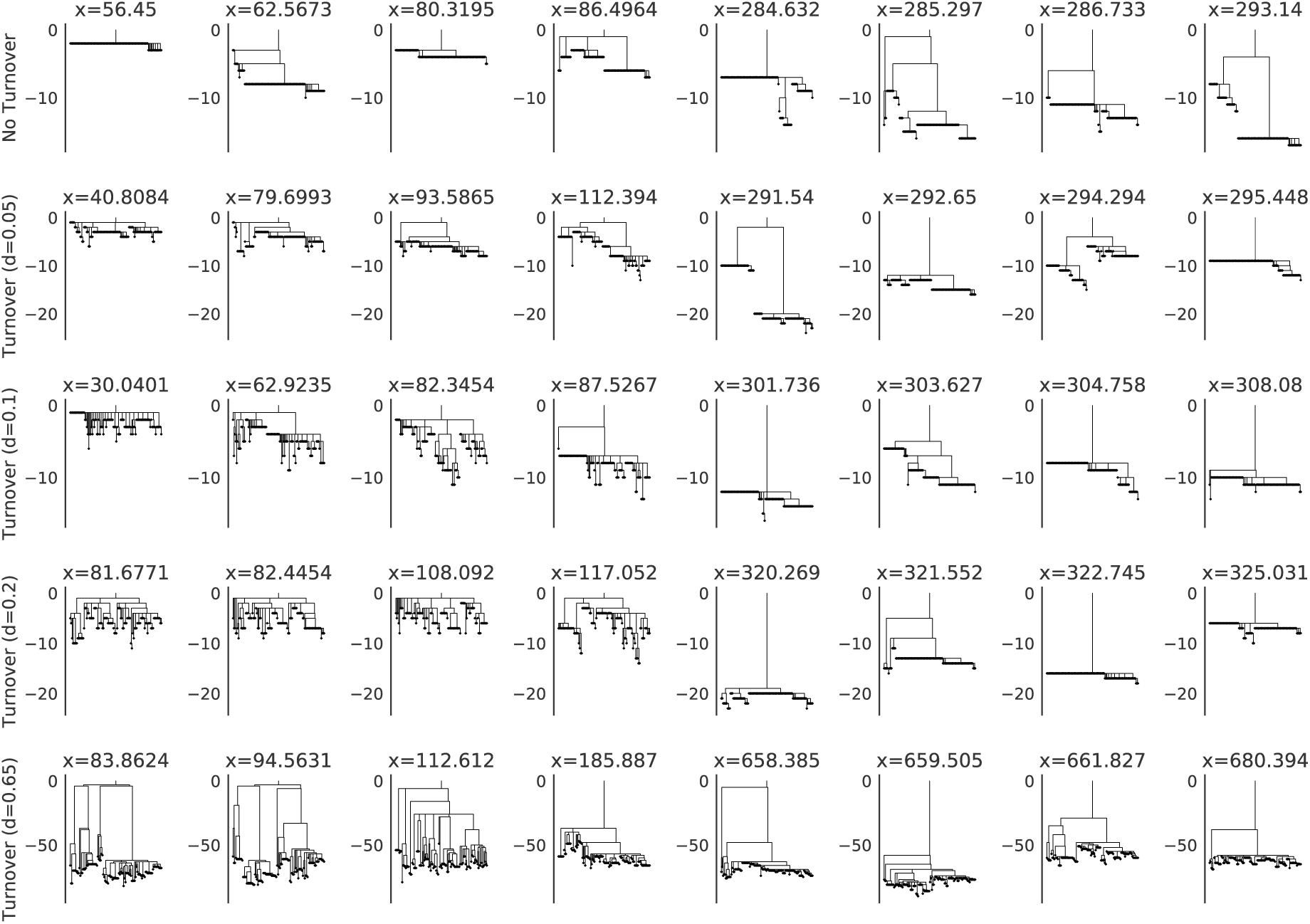
Visualizing coalescence trees for neighbourhoods in different parts of the tumor: Ancestral trees for neighbourhoods near the center (first four columns) and the edge (last four columns) for different tumor models, where branch length indicates the number of mutations that occurred on that branch. *x* is the distance from the tumor center at which the neighbourhood was sampled. Trees near the center have longer terminal branches while trees near the edge have longer stems. This pattern becomes more pronounced as the death rate is increased.

**Supplemental Figure 11:**
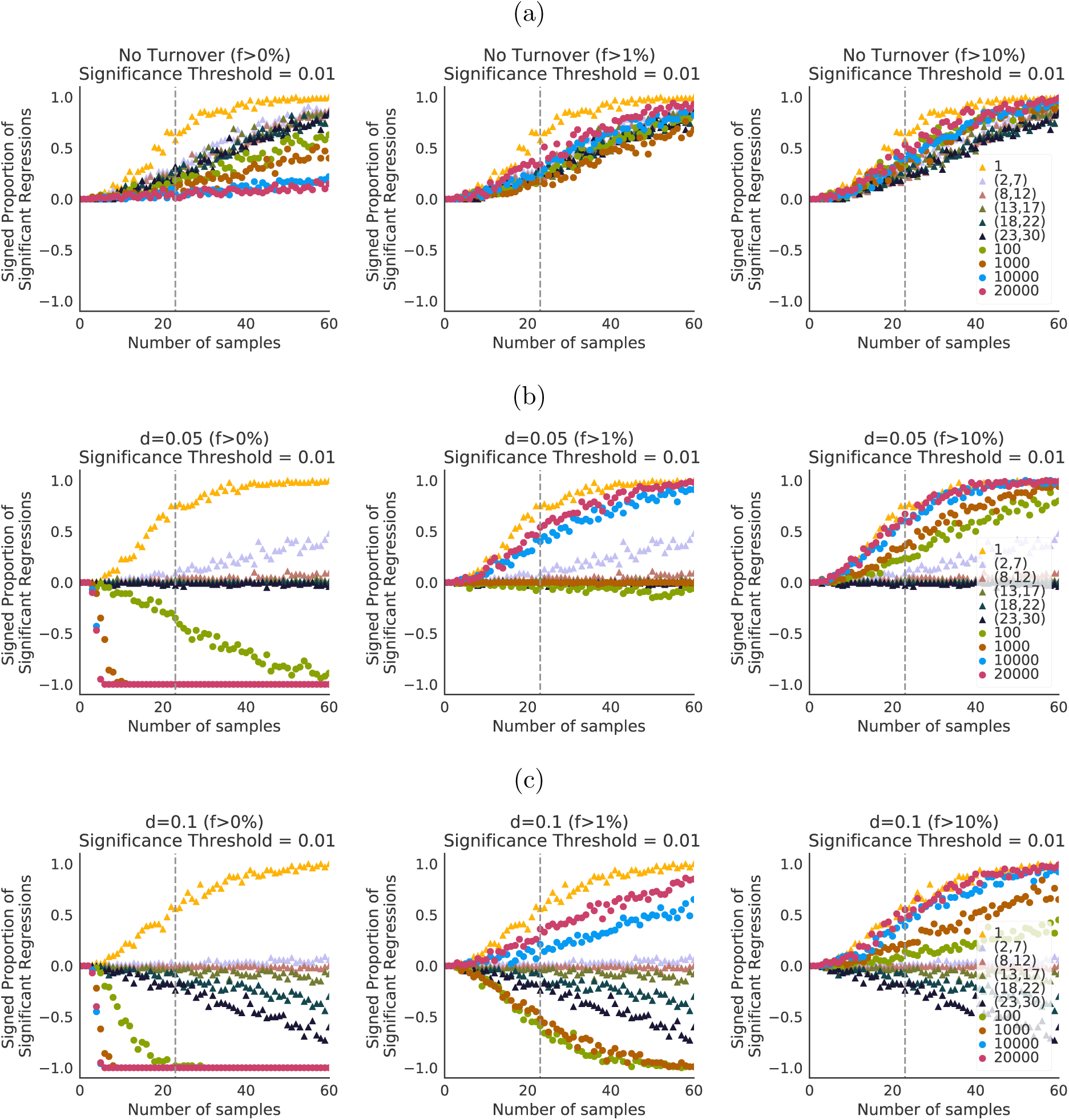
Number of samples necessary to detect spatial trends from a regression analysis for CTCs and biopsies in the models where (a) *d* = 0, (b) *d* = 0.05 and (c) *d* = 0.1. Frequency cutoff for small cell clusters is 0% (i.e., we detect all mutations), and we let cutoffs vary from 0% to 10% for large clusters (to reflect values used in dataset from Ling *et al.* (2015)). By increasing the focus on common, older mutations, the imposition of a cutoff qualitatively changes spatial trends of diversity, hiding the effect of rare, recent variants observed in Fig 2.

**Supplemental Figure 12:**
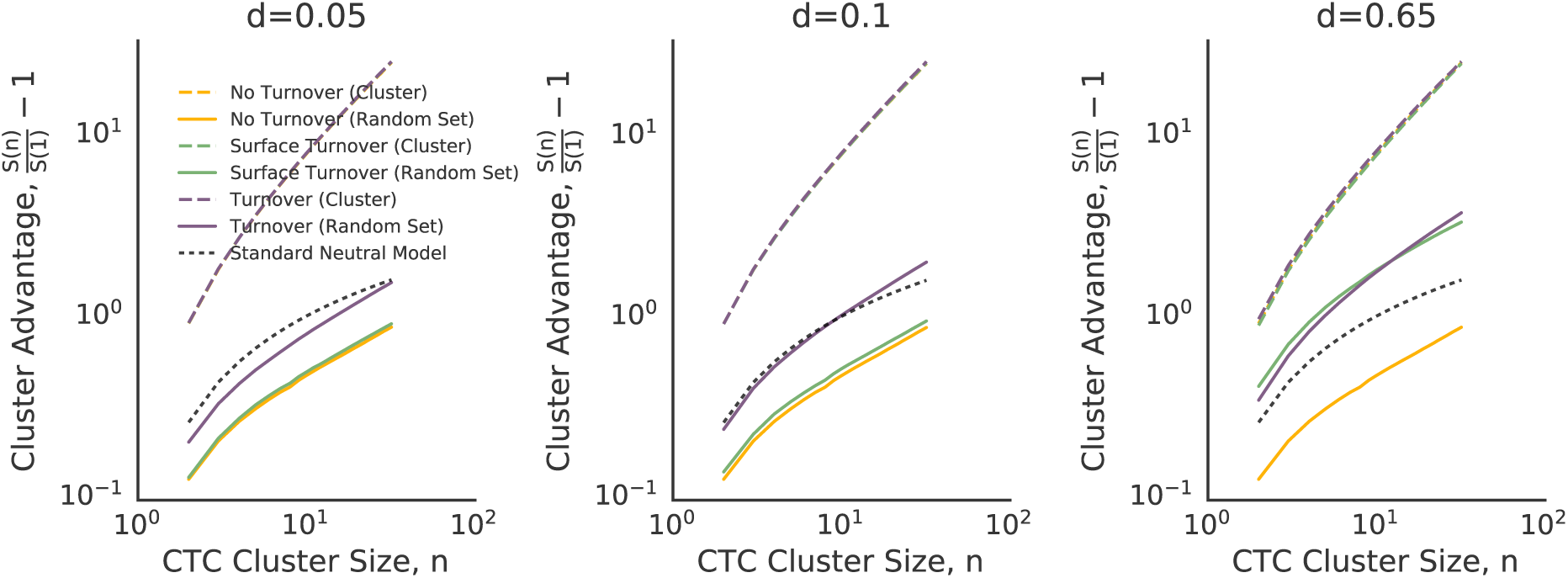
Cluster advantage for weak turnover models: even weak mixing (turnover model with *d* = 0.05) can lead to substantial differences in the cluster advantage.

### Table S2: Parameters for all reported simulations

The code to run all simulations presented here can be found at <u>this online repository</u>. These parameters must be specified in `params.h`. Alternatively, all parameters are pre-written into the repository and can be compiled in one command using `compile_all_experiments.sh`. Driver mutation rate (`driver_prob`) is fixed to 2e-5. Tumors are grown to size 10^8^, unless specified. To toggle between surface turnover models, core turnover column is specified to be either ON or OFF. If ON, the line `DEATH_ON_SURFACE` must be uncommented in the params file. Initial birth rate is specified as `growth0` and is set to 0.69. Parameters for individual simulations are reported below.

### Passenger Mutation Rate 1e-2

| Figure | Experiment Name | Death Rate<br>(`death0`) | Selection Coefficient<br>(`driver_adv`) | Core Turnover<br>(`DEATH_ON_SURFACE`) | Executable name on<br>repository once<br>compiled |
| --- | --- | --- | --- | --- | --- |
| Fig 1 Frequency Spectra | d=0.05, no turnover | 0 | 0.01 | OFF | 1_0_0 |
|  | d=0.05, surface turnover | 0.05 | 0.01 | ON | 1_1_005 |
|  | d=0.05, turnover | 0.05 | 0.01 | OFF | 1_0_01 |
|  | d=0.65, no turnover | 0 | 0.01 | OFF | 1_0_0 |
|  | d=0.65, surface turnover | 0.65 | 0.01 | ON | 1_1_065 |
|  | d=0.65, turnover | 0.65 | 0.01 | OFF | 1_0_065 |
| Fig S1 Frequency Spectra | d=0.1, no turnover | 0 | 0.01 | OFF | 1_0_0 |

“Intratumor Heterogeneity and Circulating Tumor Cell Clusters” (Ahmed and Gravel, 2018)

|  |  |  |  |  |  |
| --- | --- | --- | --- | --- | --- |
|  | d=0.1, surface turnover | 0.1 | 0.01 | ON | 1_1_01 |
|  | d=0.1, turnover | 0.1 | 0.01 | OFF | 1_0_01 |
|  | d=0.2, no turnover | 0 | 0.01 | OFF | 1_0_0 |
|  | d=0.2, surface turnover | 0.2 | 0.01 | ON | 1_1_02 |
|  | d=0.2, turnover | 0.2 | 0.01 | OFF | 1_0_02 |
| Fig S2(a), Frequency spectra vs selection rate | No Selection, d=0.0 | 0 | 0.0 | OFF | 0_0_0 |
|  | No Selection, d=0.05 | 0.05 | 0.0 | ON | 0_0_005 |
|  | No Selection, d=0.1 | 0.1 | 0.0 | OFF | 0_0_01 |
|  | No Selection, d=0.2 | 0.2 | 0.0 | OFF | 0_0_02 |
|  | No Selection, d=0.65 | 0.65 | 0.0 | ON | 0_0_065 |
| Fig S2(c), frequency spectra vs selection rate | Selection 10%, d=0.0 | 0 | 0.1 | OFF | 10_0_0 |
|  | Selection 10%, d=0.05 | 0.05 | 0.1 | ON | 10_0_005 |
|  | Selection 10%, d=0.1 | 0.1 | 0.1 | OFF | 10_0_01 |
|  | Selection 10%, d=0.2 | 0.2 | 0.1 | OFF | 10_0_02 |
|  | Selection 10%, d=0.65 | 0.65 | 0.1 | ON | 10_0_065 |

### Passenger Mutation Rate 1.875e-2

| Figure | Experiment Name | Death Rate (death0) | Selection Coefficient (driver_adv) | Core Turnover (DEATH_ON_SURFACE) | Executable name in repository once compiled |
| --- | --- | --- | --- | --- | --- |
| Fig 2 Number of somatic mutations per cluster | No Turnover | 0 | 0.01 | OFF | 1_0_0 |
|  | Turnover, d=0.05 | 0.05 | 0.01 | OFF | 1_0_005 |
|  | Turnover, d=0.1 | 0.1 | 0.01 | OFF | 1_0_01 |
|  | Turnover d=0.2 | 0.2 | 0.01 | OFF | 1_0_02 |
| Fig S4 | Turnover d=0.2 | 0.65 | 0.01 | OFF | 1_0_065 |
| Fig 2 Number of somatic mutations per cluster | Surface Turnover, d=0.05 | 0.05 | 0.01 | ON | 1_1_005 |
|  | Surface Turnover, d=0.1 | 0.1 | 0.01 | ON | 1_1_01 |
|  | Surface Turnover d=0.2 | 0.2 | 0.01 | ON | 1_1_02 |
|  | Surface Turnover, d=0.65 | 0.65 | 0.01 | ON | 1_1_065 |

## Simulation Compilation and Submission

The script <u>compile_all_experiments.sh</u> will compile all the experiments according to the above parameters. If you are on a cluster you can use <u>submit_all_experiments.sh</u> to submit all of them to a queue. This script is called multiple times with different mutation rates u0.01 and u0.01875 and seeds: `[′10′,′100′,′102′,′15′,′3′,′3318′,′33181′,′33185′,′33186′,′34201810′,′342018101′,′342018102′,′8′,′9′,′9 9′]`

### Analysis Pipeline

See https://github.com/zafarali/tumorheterogeneity/blob/mixing-parallel/analysis/_init_py

| Code Module | Purpose |
| --- | --- |
| load_tumor | Loads the tumor into memory |
| create_kdsampler | Creates the sampler to search for SNPs in the tumor |
| marginal_counts_unordered | Used for the advantage plots with random sampling |
| marginal_counts_ordered | Used for the advantage plots with ordered sampling |
| density_plot | Density plot in the background |
| big_samples | Gets the big samples from the tumor (upwards of 10k) |
| perform_mixing_analysis | Performs mixing analysis |

### Pipeline for figures

See <u>run_figure_pipeline.sh</u>

~~~
PATH_TO_ALL_SIMS=″../model/experiments/u0.01875/″
SELECTED_TUMOR=″10″ # seed of the selected tumor for analysis

python2 do_power_analysis.py $PATH_TO_ALL_SIMS $SELECTED_TUMOR
python2 create_AFS_figures.py
python2 create_number_of_divisions_table.py > S1table.txt
python2 create_Splots_figures.py $SELECTED_TUMOR
python2 create_cluster_advantage_plots.py
python2 create_fanning_plots_v2.py
python2 create_trees.py $PATH_TO_ALL_SIMS $SELECTED_TUMOR
python2 create_analytic_plot.py
~~~

### Recreating Experiments in S7 and S8

Switch to the branch `TRANSITION_EXPERIMENT` and run the compile script <u>compile_transition_experiments.sh</u>

You can then use <u>submit_all_experiments.sh</u> (`bash submit_all_experiments.sh u0.01transition` seed dry) to submit them to a cluster. The seeds used for this experiment are [6, 7, 8]

### Recreating Experiments in S4

Switch to branch `new-death-models` and run the compile script. You can then use <u>submit_all_experiments.sh</u> (bash submit_all_experiments.sh u0..01lowcutoff SEED DRY) to submit them to a cluster. The seeds used for this experiment are [1, 2, 3, 4, 5]

“Intratumor Heterogeneity and Circulating Tumor Cell Clusters” (Ahmed and Gravel, 2018)

## Footnotes

1 https://github.com/zafarali/tumorheterogeneity

2 http://www2.ph.ed.ac.uk/ bwaclaw/cancer-code/

## References

Aceto, N., A. Bardia, et al. (2014). “Circulating tumor cell clusters are oligoclonal precursors of breast cancer metastasis”. Cell 158.5, 1110–1122.

Aceto, N., M. Toner, et al. (2015). “En route to metastasis: circulating tumor cell clusters and epithelial-to-mesenchymal transition”. Trends in Cancer 1.1, 44–52.

Alizadeh, A. A. et al. (2015). “Toward understanding and exploiting tumor heterogeneity”. Nature Medicine 21.8, 846–853.

Andor, N. et al. (2016). “Pan-cancer analysis of the extent and consequences of intratumor heterogeneity”. Nature Medicine 22.1, 105.

Ashworth, T. (1869). “A case of cancer in which cells similar to those in the tumours were seen in the blood after death”. Aust Med J. 14, 146.

Au, S. H. et al. (2017). “Microfluidic isolation of circulating tumor cell clusters by size and asymmetry”. Scientific Reports 7.1, 2433.

Bos, P. D. et al. (2009). “Genes that mediate breast cancer metastasis to the brain”. Nature 459.7249, 1005–1009.

Brouwer, A. et al. (2016). “Evaluation and consequences of heterogeneity in the circulating tumor cell compartment”. Oncotarget 7.30, 48625.

Burrell, R. A. et al. (2013). “The causes and consequences of genetic heterogeneity in cancer evolution”. Nature 501.7467, 338.

Cheung, K. J. et al. (2016). “Polyclonal breast cancer metastases arise from collective dissemination of keratin 14-expressing tumor cell clusters”. Proceedings of the National Academy of Sciences 113.7, E854–E863.

Cristofanilli, M. et al. (2005). “Circulating tumor cells: a novel prognostic factor for newly diagnosed metastatic breast cancer”. Journal of Clinical Oncology 23.7, 1420–1430.

Del Monte, U. (2009). “Does the cell number 10^9^ still really fit one gram of tumor tissue?” Cell Cycle 8.3, 505–506.

Durrett, R. (2008). Probability models for DNA sequence evolution. Springer Science & Business Media.

Enriquez-Navas, P. M. et al. (2016). “Exploiting evolutionary principles to prolong tumor control in preclinical models of breast cancer”. Science Translational Medicine 8.327, 327ra24–327ra24.

Excoffier, L., M. Foll, and R. J. Petit (2009). “Genetic consequences of range expansions”. Annual Review of Ecology, Evolution, and Systematics 40, 481–501.

Fisher, R. A. (1999). The genetical theory of natural selection: a complete variorum edition. Oxford University Press.

Fusco, D. et al. (2016). “Excess of mutational jackpot events in expanding populations revealed by spatial Luria–Delbrück experiments”. Nature Communications 7, 12760.

Gerlinger, M., S. Horswell, et al. (2014). “Genomic architecture and evolution of clear cell renal cell carcinomas defined by multiregion sequencing”. Nature Genetics 46.3, 225–233.

Gerlinger, M., A. J. Rowan, et al. (2012). “Intratumor heterogeneity and branched evolution revealed by multiregion sequencing”. New England Journal of Medicine 2012.366, 883–892.

Glaves, D. (1983). “Correlation between circulating cancer cells and incidence of metastases”. British Journal of Cancer 48.5, 665.

Glynn, M. et al. (2015). “Cluster size distribution of cancer cells in blood using stopped-flow centrifugation along scale-matched gaps of a radially inclined rail”. Microsystems & Nano-engineering 1, 15018.

Hallatschek, O. et al. (2007). “Genetic drift at expanding frontiers promotes gene segregation”. Proceedings of the National Academy of Sciences 104.50, 19926–19930.

Hao, J.-J. et al. (2016). “Spatial intratumoral heterogeneity and temporal clonal evolution in esophageal squamous cell carcinoma”. Nature Genetics 48.12, 1500.

Hayes, D. et al. (2002). “Monitoring expression of HER-2 on circulating epithelial cells in patients with advanced breast cancer”. International Journal of Oncology 21.5, 1111–1117.

Heitzer, E. et al. (2013). “Complex tumor genomes inferred from single circulating tumor cells by array-CGH and next-generation sequencing”. Cancer Research 73.10, 2965–2975.

Hiley, C. et al. (2014). “Deciphering intratumor heterogeneity and temporal acquisition of driver events to refine precision medicine”. Genome Biology 15.8, 453.

Hodgkinson, C. L. et al. (2014). “Tumorigenicity and genetic profiling of circulating tumor cells in small-cell lung cancer”. Nature Medicine 20.8, 897–903.

Holohan, C. et al. (2013). “Cancer drug resistance: an evolving paradigm”. Nature Reviews Cancer 13.10, 714–726.

Hou, J. M. et al. (2012). “Clinical significance and molecular characteristics of circulating tumor cells and circulating tumor microemboli in patients with small-cell lung cancer”. Journal of Clinical Oncology 30.5, 525–532.

Jamal-Hanjani, M., A. Hackshaw, et al. (2014). “Tracking genomic cancer evolution for precision medicine: the lung TRACERx study”. PLoS Biology 12.7, e1001906.

Jamal-Hanjani, M., G. A. Wilson, et al. (2017). “Tracking the evolution of non–small-cell lung cancer”. New England Journal of Medicine 376.22, 2109–2121.

Joosse, S. A., T. M. Gorges, and K. Pantel (2014). “Biology, detection, and clinical implications of circulating tumor cells”. EMBO Molecular Medicine, e201303698.

Jouganous, J. et al. (2017). “Inferring the joint demographic history of multiple populations: beyond the diffusion approximation”. Genetics, 117.

Kardar, M., G. Parisi, and Y.-C. Zhang (1986). “Dynamic scaling of growing interfaces”. Physical Review Letters 56.9, 889.

Korolev, K. S. et al. (2010). “Genetic demixing and evolution in linear stepping stone models”. Reviews of Modern Physics 82.2, 1691.

Krebs, M. G., R. L. Metcalf, et al. (2014). “Molecular analysis of circulating tumour cells-biology and biomarkers.” Nature Reviews Clinical Oncology 11.3, 129–44.

Krebs, M. G., R. Sloane, et al. (2011). “Evaluation and prognostic significance of circulating tumor cells in patients with non–small-cell lung cancer”. Journal of Clinical Oncology 29.12, 1556–1563.

Lambert, A. W., D. R. Pattabiraman, and R. A. Weinberg (2017). “Emerging biological principles of metastasis”. Cell 168.4, 670–691.

Ling, S. et al. (2015). “Extremely high genetic diversity in a single tumor points to prevalence of non-Darwinian cell evolution”. Proceedings of the National Academy of Sciences 112.47.

Liotta, L. A., J. Kleinerman, and G. M. Saldel (1976). “The significance of hematogenous tumor cell clumps in the metastatic process”. Cancer research 36.3, 889–894.

Lorusso, G. and C. Rüegg (2012). “New insights into the mechanisms of organ-specific breast cancer metastasis”. Seminars in Cancer Biology. Vol. 22. 3. Elsevier, 226–233.

Lyons, R., R. Pemantle, and Y. Peres (1995). “Conceptual proofs of L log L criteria for mean behavior of branching processes”. The Annals of Probability, 1125–1138.

Marrinucci, D. et al. (2012). “Fluid biopsy in patients with metastatic prostate, pancreatic and breast cancers”. Physical Biology 9.1, 016003.

Massagué, J. and A. C. Obenauf (2016). “Metastatic colonization by circulating tumour cells”. Nature 529.7586, 298–306.

McGranahan, N. and C. Swanton (2015). “Biological and therapeutic impact of intratumor heterogeneity in cancer evolution”. Cancer Cell 27.1, 15–26.

McGranahan, N. and C. Swanton (2017). “Clonal heterogeneity and tumor evolution: past, present, and the future”. Cell 168.4, 613–628.

Morrissy, A. S. et al. (2017). “Spatial heterogeneity in medulloblastoma”. Nature Genetics 49.5, 780.

Navin, N. et al. (2010). “Inferring tumor progression from genomic heterogeneity”. Genome Research 20.1, 68–80.

Nowell, P. C. (1976). “The clonal evolution of tumor cell populations”. Science 194.4260, 23–28.

Ohtsuki, H. and H. Innan (2017). “Forward and backward evolutionary processes and allele frequency spectrum in a cancer cell population”. Theoretical Population Biology 117, 43–50.

Padua, D. et al. (2008). “TGFβ primes breast tumors for lung metastasis seeding through angiopoietin-like 4”. Cell 133.1, 66–77.

Peinado, H. et al. (2017). “Pre-metastatic niches: organ-specific homes for metastases”. Nature Reviews Cancer 17.5, 302.

Powell, A. A. et al. (2012). “Single cell profiling of circulating tumor cells: transcriptional heterogeneity and diversity from breast cancer cell lines”. PloS ONE 7.5, e33788.

Quail, D. F. and J. A. Joyce (2013). “Microenvironmental regulation of tumor progression and metastasis”. Nature Medicine 19.11, 1423.

Sarioglu, A. F. et al. (2015). “A microfluidic device for label-free, physical capture of circulating tumor cell clusters”. Nature Methods 12.7, 685.

Schneider, C. A., W. S. Rasband, and K. W. Eliceiri (2012). “NIH Image to ImageJ: 25 years of image analysis”. Nature Methods 9.7, 671.

Shweiki, D. et al. (1995). “Induction of vascular endothelial growth factor expression by hypoxia and by glucose deficiency in multicell spheroids: implications for tumor angiogenesis”. Proceedings of the National Academy of Sciences 92.3, 768–772.

Siravegna, G. et al. (2017). “Integrating liquid biopsies into the management of cancer”. Nature Reviews Clinical Oncology 14.9, 531.

Sottoriva, A. et al. (2015). “A Big Bang model of human colorectal tumor growth”. Nature Genetics 47.3, 209–216.

Steeg, P. S. (2016). “Targeting metastasis”. Nature Reviews Cancer 16.4, 201.

Vanharanta, S. and J. Massagué (2013). “Origins of metastatic traits”. Cancer Cell 24.4, 410–421.

Waclaw, B. et al. (2015). “A spatial model predicts that dispersal and cell turnover limit intratumour heterogeneity”. Nature 525.7568, 261–264.

Wang, Y. et al. (2014). “Clonal evolution in breast cancer revealed by single nucleus genome sequencing”. Nature 512.7513, 155–160.

Weinstein, B. T. et al. (2017). “Genetic drift and selection in many-allele range expansions”. PLoS Computational Biology 13.12, e1005866.

Williams, M. J. et al. (2016). “Identification of neutral tumor evolution across cancer types”. Nature Genetics 48, 238–244.

Wright, S. (1931). “Evolution in Mendelian populations”. Genetics 16.2, 97–159.

Wülfing, P. et al. (2006). “HER2-positive circulating tumor cells indicate poor clinical outcome in stage I to III breast cancer patients”. Clinical Cancer Research 12.6, 1715–1720.

Yates, L. R. et al. (2015). “Subclonal diversification of primary breast cancer revealed by multiregion sequencing”. Nature Medicine 21.7, 751–759.

Zhang, J. et al. (2014). “Intratumor heterogeneity in localized lung adenocarcinomas delineated by multiregion sequencing”. Science 346.6206, 256–259.

